# Rab11B is required for binding and entry of recent H3N2, but not H1N1, influenza A isolates

**DOI:** 10.1101/2025.04.23.650275

**Authors:** Allyson H. Turner, Sara A. Jaffrani, Hannah C. Kubinski, Deborah P. Ajayi, Matthew B. Owens, Conor D. Fanuele, Madeline P. McTigue, Cailey L. Appenzeller, Addington Bowling, Hannah W. Despres, Madaline M. Schmidt, Dave J. Shirley, Jessica W. Crothers, Ramiro Barrantes-Reynolds, Emily A. Bruce

**Affiliations:** Department of Microbiology and Molecular Genetics, University of Vermont, Burlington, Vermont, USA; Cellular, Molecular, and Biomedical Sciences Graduate Program, University of Vermont Larner College of Medicine, Burlington, VT, USA; Faraday, Inc. Data Science Department, Burlington, Vermont, United States of America; Department of Pathology and Laboratory Medicine, University of Vermont, Burlington, VT, USA

## Abstract

As an obligate intracellular parasite, influenza A virus (IAV) depends on host proteins to complete several important functions, including trafficking viral proteins throughout the cell. Previous work has established a critical role for the cellular vesicular trafficking protein, Rab11A, in transporting the viral genome segments to the site of budding at the plasma membrane. While the role of Rab11A in IAV assembly is relatively well understood, very little is known about the function of a closely related isoform (Rab11B) during influenza virus infection. We have shown that both Rab11A and Rab11B are required for successful IAV infection by current H1N1 or H3N2 isolates. Cells in which either Rab11A or Rab11B were depleted failed to efficiently produce virus, with significant reductions in infectious titer. Surprisingly, our data revealed that recent (2022) H3N2, but not H1N1, isolates failed to efficiently produce viral proteins in single-cycle infections when Rab11B (but not Rab11A) was depleted. Flow cytometry analysis suggests that the defect in protein production is driven by a reduction in the total number of infected cells, rather than a decrease in viral protein production at the single cell level. Using reverse genetics and ‘7+1’ reassortant viruses we mapped this Rab11B-dependent early defect in recent H3N2 isolates to the HA gene. RT-qPCR analysis of H3N2 virions bound to the cell surface showed a ∼50% decrease in virus binding the surface of cells depleted of Rab11B, but not Rab11A. This data suggests a novel role for Rab11B during viral entry, likely at the stage of viral binding, that is specific to H3N2 isolates.

## INTRODUCTION

Influenza A virus (IAV) is a highly infectious pathogen, causing seasonal epidemics that result in around three to five million severe cases each year, in addition to causing periodic pandemics that emerge from novel IAV strains^1,2^. While there are many different subtypes of IAV, currently there are only two which are circulating within the human population, H1N1 and H3N2^3^. Influenza owes much of its pandemic potential to the fact that its genome is composed of eight distinct segments which can mix to form novel genetic combinations in a cell infected with two influenza viruses simultaneously, a process known as reassortment^4^.

As a segmented negative strand RNA virus, influenza encodes at least 14 proteins on its eight segments^3^. After transcription and translation, these proteins collectively carry out the functions required to complete the viral life cycle within a host cell. Cellular entry of the influenza virion is mediated by the viral glycoprotein hemagglutinin (HA) binding to sialic acid moieties on the cell surface, which is thought to trigger receptor mediated endocytosis^5,6^. Broadly speaking, influenza viruses that infect human cells are thought to bind to α2,6 sialic acid modifications on cell surface proteins, differentiating them from avian viruses which are thought to prefer α2,3 sialic acids [reviewed in^3,4,7^]. The classical model of influenza binding continues to be refined and expanded however, through new work that suggests there may be subtype-specific effects and both the identity of the sialylated protein and its cell surface density may play important roles^8–15^. Specific characteristics of the sialic acid modifications itself can also affect binding, with more recent H3N2 isolates preferring a highly branched structure^16^. Intriguingly H3N2s have shown decreased avidity for α2,6 sialic acid from 1968-2010 which correlates with a decrease in infection severity and evidence that NA contributed to mediating receptor binding^17^. After viral binding, internalization occurs through an incompletely understood process that is likely to involve signaling from one or more cell surface proteins and may require specific ‘internalization receptors’ [reviewed in^18,19^].

After endocytosis, the lowered pH of the endosome triggers a conformational change in HA which mediates fusion between viral and cellular membranes^20–23^. The acidic environment of the endosome also triggers activation of the matrix 2 protein (M2) ion channel, which transports protons to the interior of the virion, resulting in disassociation of M1 from the ribonucleoprotein complexes (RNPs) that is required for their subsequent nuclear import^24^. Once released into the cytoplasm, the RNP complexes that comprise the genome segments are transported to the nucleus where influenza genome replication occurs. During late stages of viral infection newly produced genome segments (vRNPS) comprising full length negative sense RNA molecules are coated with nucleoprotein (NP) and bound to the polymerase complex composed of the polymerase basic protein 1 (PB1), polymerase basic protein 2 (PB2) and the polymerase acidic protein (PA) and then exported from the nucleus. Once in the cytoplasm, vRNPs must traffic to the site of budding at the plasma membrane before being released in newly formed virions to start the cycle of infection anew^25^.

The process of transporting the vRNPs during this stage is highly dependent on the cellular Rab11 pathway, which is co-opted during viral infection to coordinate the apical transport of the influenza genome^26–28^. Rab11 is a member of the Rab GTPase family, and plays a key role in vesicular transport within the recycling endosome pathway^29^. A direct interaction between Rab11A and PB2 (a component of the influenza tripartite polymerase complex) mediates binding between Rab11A and negative sense vRNPs, with no binding observed to full length positive sense or viral mRNA species^26,30^. This interaction is hypothesized to disrupt the binding of Rab11-Family Interacting Proteins (FIPs), possibly through Rab11’s regulatory Switch I region^30^. Influenza infection disrupts the normal functioning of the endosomal trafficking pathway to favor transport of vRNPs to the plasma membrane^31–33^. During transport, large clusters of multiple genome segments that colocalize with Rab11 can be observed by fluorescent *in situ* hybridization^26^. This Rab11 dependent transport serves to concentrate and cluster the eight IAV segments in defined subcellular sites, dependent on microtubules^34–39^. Finally, Rab11 is known to play a role in the very late stages of the viral life cycle. Rab11 can traffic vRNPs between cells through tunneling nanotubules, allowing for cell to cell spread without generation of complete virions^40^. Cells lacking Rab11 fail to produce filamentous virions and exhibit signs of budding defects, possibly due to a secondary failure to transport the viral genome to the site of budding^41^.

Notably, the majority of this research has been focused exclusively on the role of Rab11A, while a small subset of studies looked at the combined effect of Rab11A and Rab11B. Studies examining the role of the different Rab11 isotypes, in the context of cancer, reveal the two isoforms can play distinct roles, leading us to wonder if there might also be differences in how they interact with viral proteins during the IAV lifecycle^42^. While the exact binding site between PB2 and Rab11A is unknown, the interaction between these proteins is completely abolished by mutations to the C-terminal Class I switch region, an area that is highly conserved between the different Rab11 family members^30^. This data, along with a previously reported drop in titer from A549 cells depleted of Rab11B and infected with PR8, suggested to us that Rab11B might play a similar role as Rab11A during influenza infection, though this phenotype did not extend to HEK-293T cells^41^. As the majority of studies examining the role of Rab11 in influenza infection have utilized older (often lab-adapted) strains, we were also curious about whether the requirement for Rab11 extended to currently circulating strains of influenza A. Here we show that both Rab11A and Rab11B are required for efficient production of infectious virions from seasonal H1N1 and H3N2 subtypes circulating in 2022. As expected, Rab11A played a role late in infection (i.e., post protein production) for both subtypes. Surprisingly, we discovered that Rab11B is required early in the life cycle of H3N2 infection, with cells lacking Rab11B failing to efficiently produce viral proteins. We were able to map this defect specifically to the HA gene and trace it to a defect early in viral entry, as H3N2 (but not H1N1) virions fail to bind to the surface of cells lacking Rab11B. Our findings suggest a surprising new role for one of the two highly conserved Rab11 isoforms in influenza infection and highlights a clear role of differential requirements for viral binding and entry of recent H1N1 vs H3N2 subtypes.

## RESULTS

To determine the effect Rab11A and Rab11B played on the replication of current seasonal influenza viruses, we used siRNAs targeting each gene to transiently knockdown expression in human adenocarcinoma alveolar basal epithelial (A549) cells, in the hopes of minimizing off-target effects that can occur with stable knockdown/knockout. In addition to Rab11A and Rab11B, we included a third Rab (Rab8A) that is also involved in recycling endosome trafficking but is not known to impact influenza viral replication, as well as a non-targeting control. To verify the efficiency of our knockdown, we initially used western blot to measure Rab11 protein expression, relative to a house keeping gene (β-actin). We observed global Rab11 protein depletion using a non-isoform specific antibody in cells treated with siRNA targeting both Rab11A and Rab11B (Figure 1A). Using a Rab11A isoform specific antibody, we could see depletion of Rab11A in cells treated with siRNA to Rab11A (or Rab11A and B in combination) (Figure 1B). However, due to difficulties in antibody sensitivity when detecting endogenous protein levels we used RT-qPCR to quantify gene expression, relative to a house-keeping transcript (18S) and expression in a NT control at 48 hrs post transfection. Our RT-qPCR results (Figure 1C) verified Rab11A gene expression was reduced to about 15% of endogenous levels, Rab11B to ∼5% and Rab8 to ∼20%. For cells in which we simultaneously depleted Rab11A and Rab11B, we used the same total amount of RNA (i.e., half the level/gene for a single knockdown) to avoid cell toxicity. In this case, depletions were slightly less efficient, with Rab11A gene expression reduced to ∼20% and Rab11B to ∼10% (Figure 1C).

**Figure 1.**
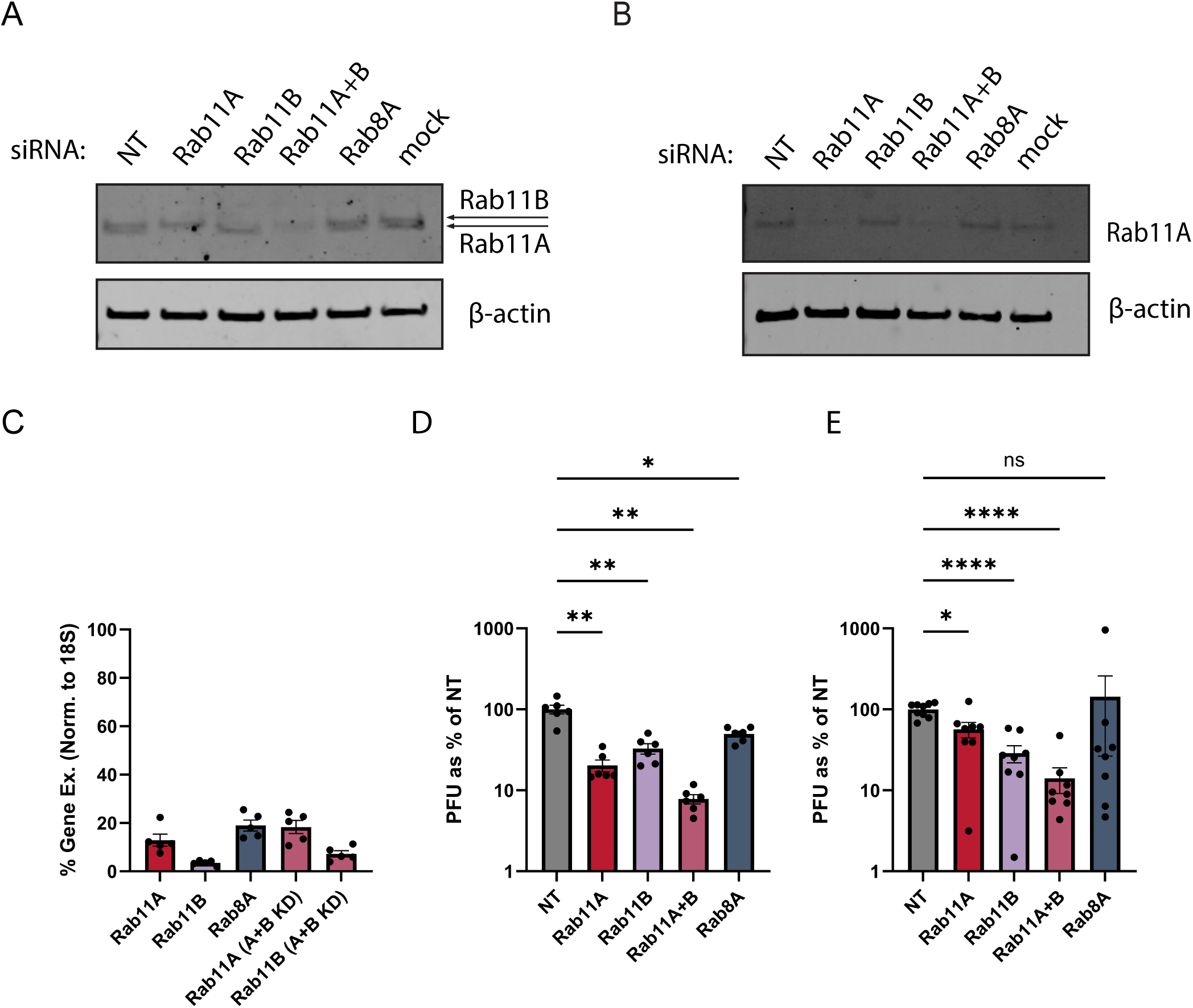
Rab11A and Rab11B are required for replication of recent H1N1 and H3N2 influenza A isolates. A549 cells were treated with siRNAs targeting Rab11A, Rab11B (singly and in combination), Rab8A or a non-targeting control. 48 hours post transfection cells were harvested to determine protein (**A, B**) or RNA (**C**) expression. **A)** Total Rab11 (Rab11A plus Rab11B) or **B)** Rab11A expression was detected by western blot using either non-specific or Rab11A specific isoform antibodies. **C)** Relative gene expression was determined by RT-qPCR, using 18S as housekeeping gene and normalized to the NT average of each biological experiment. Alternatively, at 48 hpt cells were infected with **D)** a 2022 IAV isolate of H1N1 (A/Burlington/UVM-0478/2022) or **E)** H3N2 (A/Burlington/UVM-1927/2022) subtype at an MOI of 1. Viral supernatants were collected at 16 hpi and titered by plaque assay (PFU; plaque forming unit). Mean +/- SEM of relative PFU (normalized to the NT average of each biological experiment) is plotted. Statistical comparisons (D, E) were done using Welch’s one way ANOVA with Dunnett’s multiple comparisons, [*(p<0.05), **(p<0.01), ***(p<0.001), ****(p<0.0001)]. N=6 from three biological experiments (C), N=6 from three biological experiments (D), and N=8 from four biological experiments (E) are shown.

Having confirmed the efficiency of our depletions, we then challenged each depleted condition using two different, recent, low passage IAV isolates, A/Burlington/UVM-0478/2022 (H1N1) (denoted in this study as UVM-0478) and A/Burlington/UVM-1927/2022 (H3N2) (denoted here as UVM-1927) to perform single cycle infections. As predicted, we found that cells in which either or both Rab11A/Rab11B were depleted resulted in a significant reduction in plaque forming units (PFU) of virus produced during infection. We saw a ∼3-5 fold reduction in viral titer in cells lacking Rab11A or Rab11B and a further drop in cells lacking Rab11A+B (7-10 fold), in cells infected with either UVM-0478 (Figure 1D) or UVM-1927 (Figure 1E). In cells depleted of Rab8A and infected with UVM-0478 we saw a small but consistent drop in titer (∼2 fold) while UVM-1927 replication was unaffected (raw viral titers in Figure S1).

To verify the drop in titer observed was due to the previously characterized role of Rab11 in late stage RNP trafficking, we then examined the viral protein production of infected cells in our knockdown conditions. We observed no significant effect on viral protein expression in cells infected with UVM-0478 (Figure 2A) after densitometry quantification was used to determine the levels of HA0, NP, or M2 (Figure 2B-D) normalized to the cellular housekeeping gene, GAPDH (or alternatively, normalized to β-actin, Figure S2). Unexpectedly, we observed a significant drop in viral protein production in cells depleted of Rab11B that were infected with UVM-1927 (Figure 2E). Expression of HA0, NP, and M2 (Figure 2F-H) was reduced to 25% of that seen in cells treated with the NT control. Surprisingly, this defect in viral protein production was partially rescued in cells simultaneously depleted of Rab11A and Rab11B. In order to determine if this rescue was due to less efficient knockdown (as cells treated with siRNA to Rab11A and B simultaneously received half the total siRNA that was used in individual knockdowns), we treated cells with a half dose of Rab11A or Rab11B siRNA (along with a non-targeting control) before infecting them with UVM-1927 as above. The failure to produce viral proteins was not affected by lowering the siRNA dose, with cells lacking Rab11B once again producing 80-90% less HA, NP and M2 (Figure S3). Given prior reports of reciprocal functions in the Rab11A and Rab11B isoforms, this finding suggests that Rab11A and B could also be playing opposing functions in the context of H3N2 entry^42,43^.

**Figure 2.**
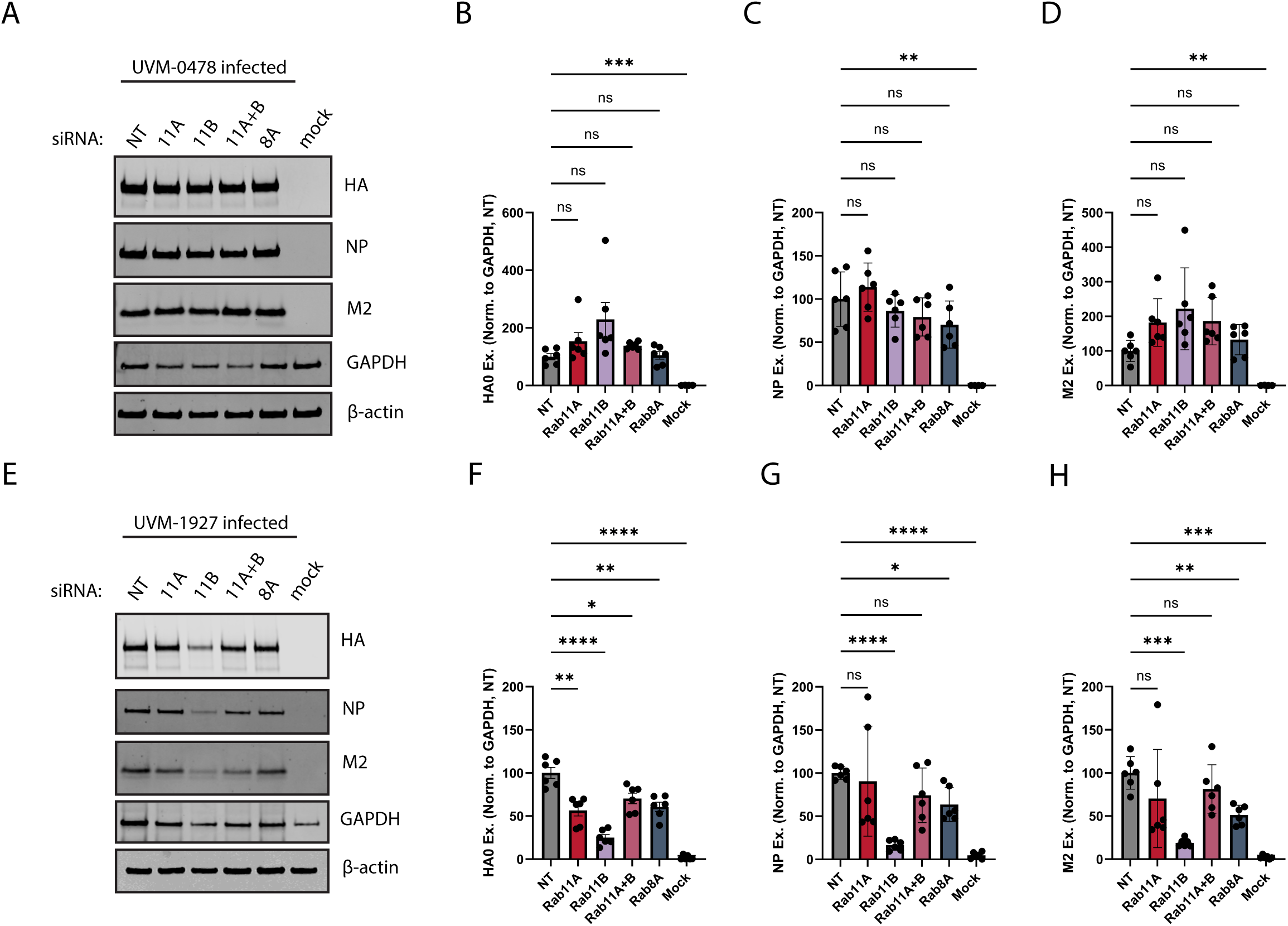
Rab11B is required early in the life cycle of a recent H3N2 IAV isolate. A549 cells were treated with siRNAs targeting Rab11A, Rab11B (singly and in combination), Rab8A or a non-targeting control, 48 hpt cells were infected with **A)** 2022 IAV isolate of H1N1 (A/Burlington/UVM-0478/2022) at an MOI of 1, or mock infected. 16 hpi cell lysates were collected and visualized by SDS-PAGE and western blot using rabbit anti-HA and anti-GAPDH antibodies in addition to mouse anti-NP and anti-M2 antibodies. Expression of **B)** HA, **C)** NP, and **D)** M2 was quantified and normalized to GAPDH levels and the NT control. Alternatively, cells were infected with **E)** 2022 IAV isolate of H3N2 (A/Burlington/UVM-1927/2022) at an MOI of 1, or mock infected. 16 hpi cell lysates were collected and visualized by SDS-PAGE and western blot using rabbit anti-HA and anti-GAPDH antibodies in addition to mouse anti-NP and anti-M2 antibodies. Expression of **F)** HA, **G)** NP, and **H)** M2 was quantified and normalized to GAPDH levels and the average NT control for each biological replicate, quantification from six technical replicates (three biological experiments) is shown. Mean +/- SEM is plotted. Statistical comparisons (B, C, D, F, G, H) were done using the Welch’s one way ANOVA with Dunnett’s multiple comparisons [*(p<0.05), **(p<0.01), ***(p<0.001), ****(p<0.0001)].

To broaden our conclusions, we obtained a second 2022 H3N2 from a different geographical region, A/Baltimore/JH-0586/2022 (H3N2) (referred to in this study as JH-0586). JH-0586 is genetically distinct from UVM-1927, with more than 100 nucleotide level differences across the eight segments of the two viruses. Like UVM-1927, JH-0586 also depended on the Rab11 isoforms to produce infectious virus, with a particular dependence on Rab11B, as observed above with UVM-1927 (Figure 3A). The loss of Rab11B resulted in a five-fold drop in viral titer while the simultaneous depletion of Rab11A and Rab11B resulted in a 20-fold decrease, while Rab8A depletion did not affect viral replication. In line with our prior observations, viral protein production was also substantially decreased in cells lacking Rab11B (Figure 3B). Quantification revealed this trend was most pronounced in expression of NP (85% of NT) and M2 (∼50%), while HA0 expression was decreased to a lesser extent (∼15%) (Figure 3C-E). Simultaneous depletion of Rab11A and Rab11B again appeared to, at least partially, rescue the defect induced by the loss of Rab11B alone. Finally, Rab8A depletion did not alter the expression of viral protein levels, compared to the NT control.

**Figure 3.**
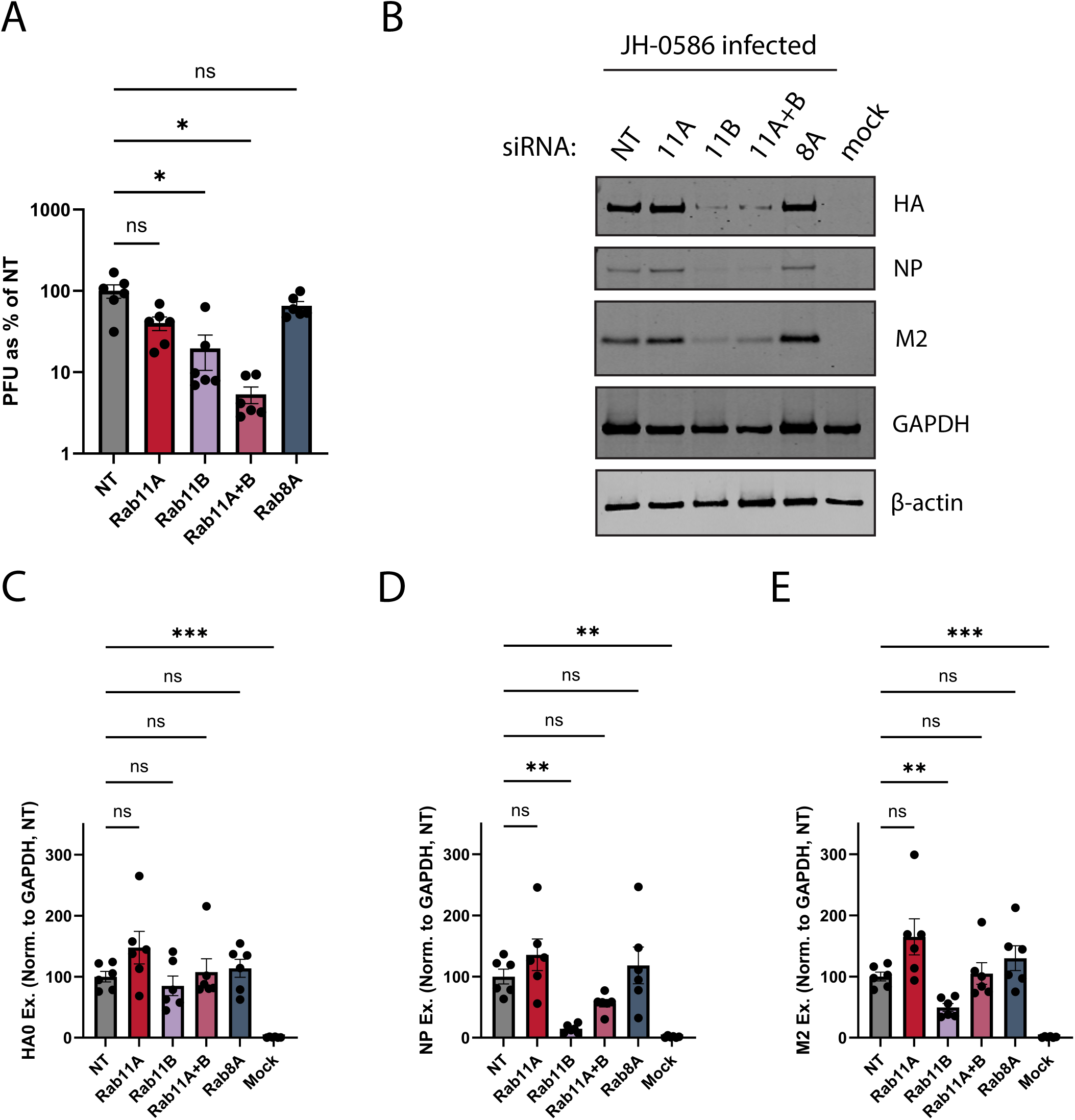
Multiple H3N2 isolates depend on Rab11B early in IAV infection. A549 cells were treated with siRNAs targeting Rab11A, Rab11B (singly and in combination), Rab8A or a non-targeting control, 48 hpt cells were infected with 2022 IAV isolate of H3N2 (A/Baltimore/JH-0586/2022) at an MOI of 1, or mock infected. **A)** 16 hpi viral supernatants were collected and titered by plaque assay (PFU; plaque forming unit). **B)** Simultaneously, cell lysates were collected and visualized by SDS-PAGE and western blot using rabbit anti-HA and anti-GAPDH antibodies in addition to mouse anti-NP and anti-M2 antibodies. Expression of viral proteins was quantified and normalized to GAPDH levels and the average of the NT controls for each biological replicate and is shown for **C)** HA0, **D)** NP and **E)** M2. Mean +/- SEM is plotted, normalized to the average of the NT control in each biological experiment N=6 from three biological experiments. Statistical comparisons (A, C, D, E) were done using Welch’s one way ANOVA with Dunnett’s multiple comparisons [*(p<0.05), **(p<0.01), ***(p<0.001)].

To determine if this phenotype was restricted to A549 cells, we next expanded our study to include NCI-H441 (H441) cells, an immortalized human ‘club cell’-like line. To rule out off-target effects of the siRNA constructs/depletion strategy we had used in A549 cells, we used lentiviral transduction to create a stable cell line with a doxycycline-driven inducible shRNA targeting Rab11B. We achieved high (∼90%) levels of Rab11B depletion in these cells three days after induction (Figure 4A), allowing us to examine viral protein production of UVM-1927 after a single round of replication (Figure 4B). Expression of HA, NP and M2 was once again significantly (<50%) decreased in cells induced with Rab11B shRNA constructs (Figure 4C-E), suggesting the dependence of recent H3N2 isolates on Rab11B was not an artifact of either cell type or our depletion strategy.

**Figure 4.**
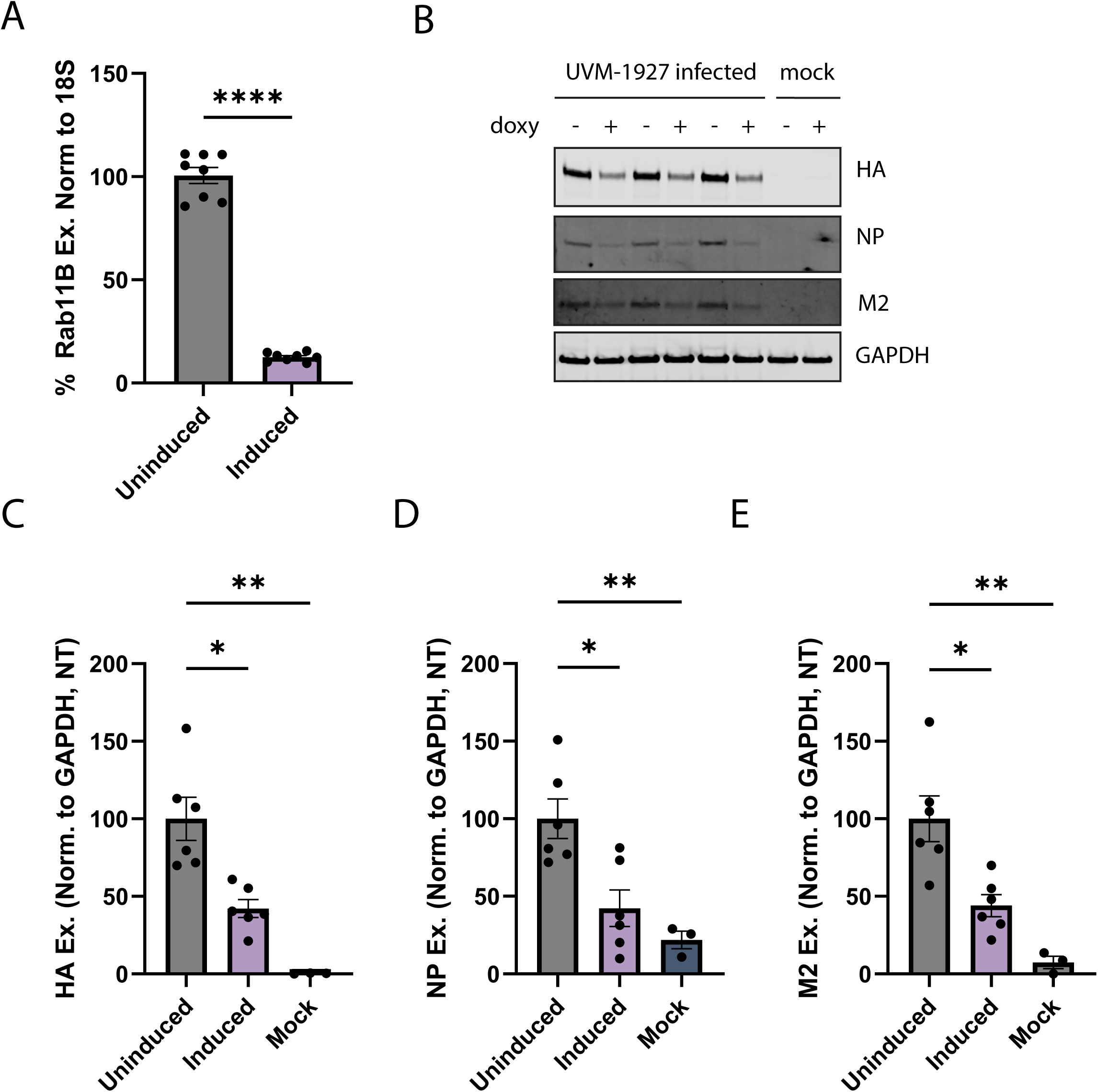
Rab11B is required early in the H3N2 life cycle in club (H441) cells. **A)** H441 cells were transduced with a lentivirus to stably express an inducible shRNA targeting Rab11B. Cells were treated (induced) or not (uninduced) with doxycycline to induce shRNA expression, and 72 hours later RNA was harvested. Rab11B gene expression was determined by RT-qPCR, using 18S as housekeeping gene and normalized to the uninduced average of each biological experiment. **B)** Alternatively, cells were infected with A/Burlington/UVM-1927/2022 or mock infected. 16 hpi cell lysates were collected and visualized by SDS-PAGE and western blot using rabbit anti-HA and anti-GAPDH antibodies in addition to mouse anti-NP and anti-M2 antibodies. Expression of **C)** HA, **D)** NP, and **E)** M2 was quantified and normalized to GAPDH levels and the average of the uninduced controls. Quantification from six technical replicates (three biological experiments) is shown. Mean +/- SEM is plotted. Statistical comparisons done using Welch’s T-test (A) and Welch’s one way ANOVA with Dunnett’s multiple comparisons (C, D, E) [*(p<0.05), **(p<0.01), ****(p<0.0001)].

Next, we wanted to understand whether our phenotype was being driven by a global drop in the number of cells being infected, or if the same number of cells were infected but each produced fewer viral proteins. As western blots of total cell lysates are unsuited to analyzing protein production at a single cell level, we used flow cytometry to measure HA and M2 expression in individual cells that had been treated with siRNAs targeting Rab11A, Rab11B or a NT control and infected with UVM-1927 as above. The infected control cells were clearly visualized as a second peak (Figure 5, light gray) with greater levels of staining than the mock infected cells (dark gray, single left-most peak) after staining with both anti-H3N2 serum and an M2 antibody (Figure 5A, B). As expected for a multiplicity of infection (MOI) of 1 (chosen due to relatively low titers of our low passage stocks), infected cells were roughly a third of the cell population. Unsurprisingly, given our prior data, the loss of Rab11A did not alter viral protein expression patterns (Figure 5A, B; compare maroon to light gray line). In cells lacking Rab11B, the total number of cells expressing viral proteins was reduced ∼50%, while we observed similar intensity of viral protein expression in the positive cell population (Figure 5A, B; compare how purple line is decreased along the *y*-axis dimension [count] but not the *x*-axis [viral antigen intensity]). Quantification of multiple biological replicates revealed a ∼50% drop in the number of cells infected (i.e., producing viral proteins) when Rab11B was knocked down, compared to both the NT control and the Rab11A knockdown (Figure 5C). Further analysis of viral protein expression within the infected cell population showed no decrease in HA or M2 expression (Figure 5D, E). This assay revealed a significant defect in the number of Rab11B depleted cells able to produce viral proteins, rather than a broad dampening of protein expression. This suggested that if a cell was able to make viral proteins at all then it would produce the normal amount, implying a defect upstream of translation.

**Figure 5.**
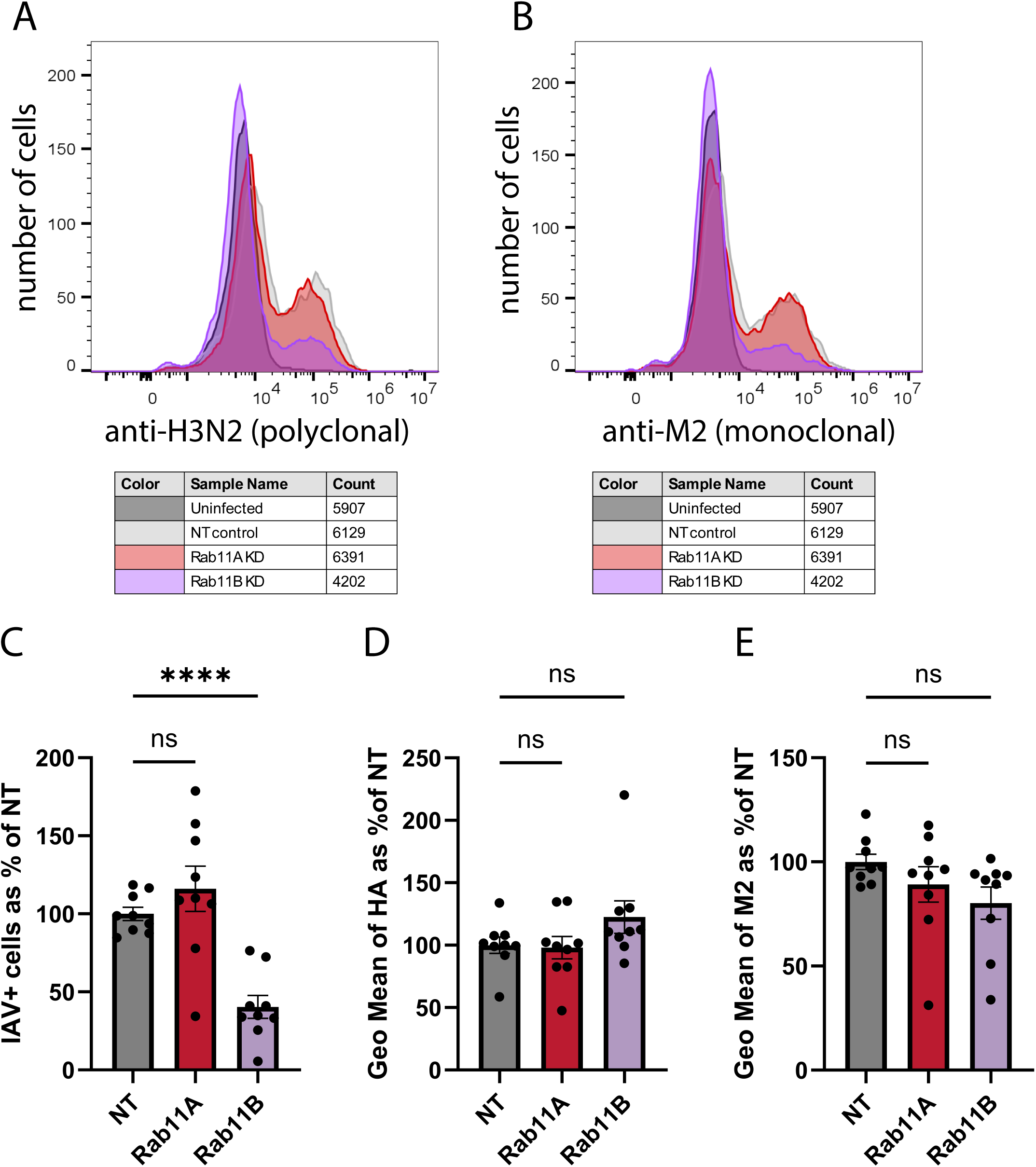
Loss of Rab11B decreases the number of H3N2 IAV infected cells rather than their protein expression. A549 cells were treated with siRNAs targeting Rab11A, Rab11B or a non-targeting control, 48 hpt cells were infected with 2022 IAV isolate of H3N2 (A/Burlington/UVM-1927/2022) at an MOI of 1, or mock infected. 16 hours post infection cells were washed with PBS, trypsinized, fixed and stained for flow cytometry analysis to determine total (intracellular/extracellular) staining using **A)** anti-H3N2 or **B)** anti-M2 antibodies. **C)** The percent of double HA^+^/M2^+^ cells in each experiment was determined using the mock control to determine background staining levels. The geographic mean of **D)** HA or **E)** M2 expression in the infected (HA^+^/M2^+^) cell population, normalized to that of the NT infected population is shown. Mean +/- SEM is plotted, normalized to the average of the NT control in each biological experiment, N=9 from three biological experiments. Statistical comparisons (C, D, E) done using Welch’s one way ANOVA with Dunnett’s multiple comparisons [****(p<0.0001)].

As our data suggested that protein (and likely RNA) production was unaffected by Rab11B depletion, we hypothesized that the infection defect in these cells was occurring somewhere in the entry process. To look at the kinetics of entry we used a ‘time of addition’ assay in which we added ammonium chloride at specific timepoints post infection to stop endosomal acidification and thus viral fusion^44^. Infections were allowed to progress in the presence of ammonium chloride, and the percentage of infected cells was measured by flow cytometry sixteen hours later (Figure 6A). In cells treated with the NT control, we saw no escape from the endosome when ammonium chloride was added concurrently with viral inoculum (Figure 6B). The majority of viral fusion appeared to occur between 20 and 60 minutes, consistent with previously reported half-times for influenza viral entry^44^. Cells lacking Rab11A showed no defect in entry kinetics, with an equal or greater number of cells infected throughout the measured time course (Figure 6B, C). In Rab11B depleted cells, we observed a significant drop in the number of infected cells at 45, 60 and 90 minutes compared to the NT control (Figure 6B, C). To better visualize whether the Rab11B entry kinetics were delayed compared to NT (which could imply utilization of an alternative trafficking pathway) we normalized the percent of infected cells to those infected in the NT control at each timepoint (Figure 6D). This revealed that cells lacking Rab11B had significant defects in their ability to support normal viral entry, but the rate of entry itself was not delayed compared to the NT control (Figure 6D), with a near constant drop in number of infected cells at each of the later timepoints compared to NT (Figure 6E).

**Figure 6.**
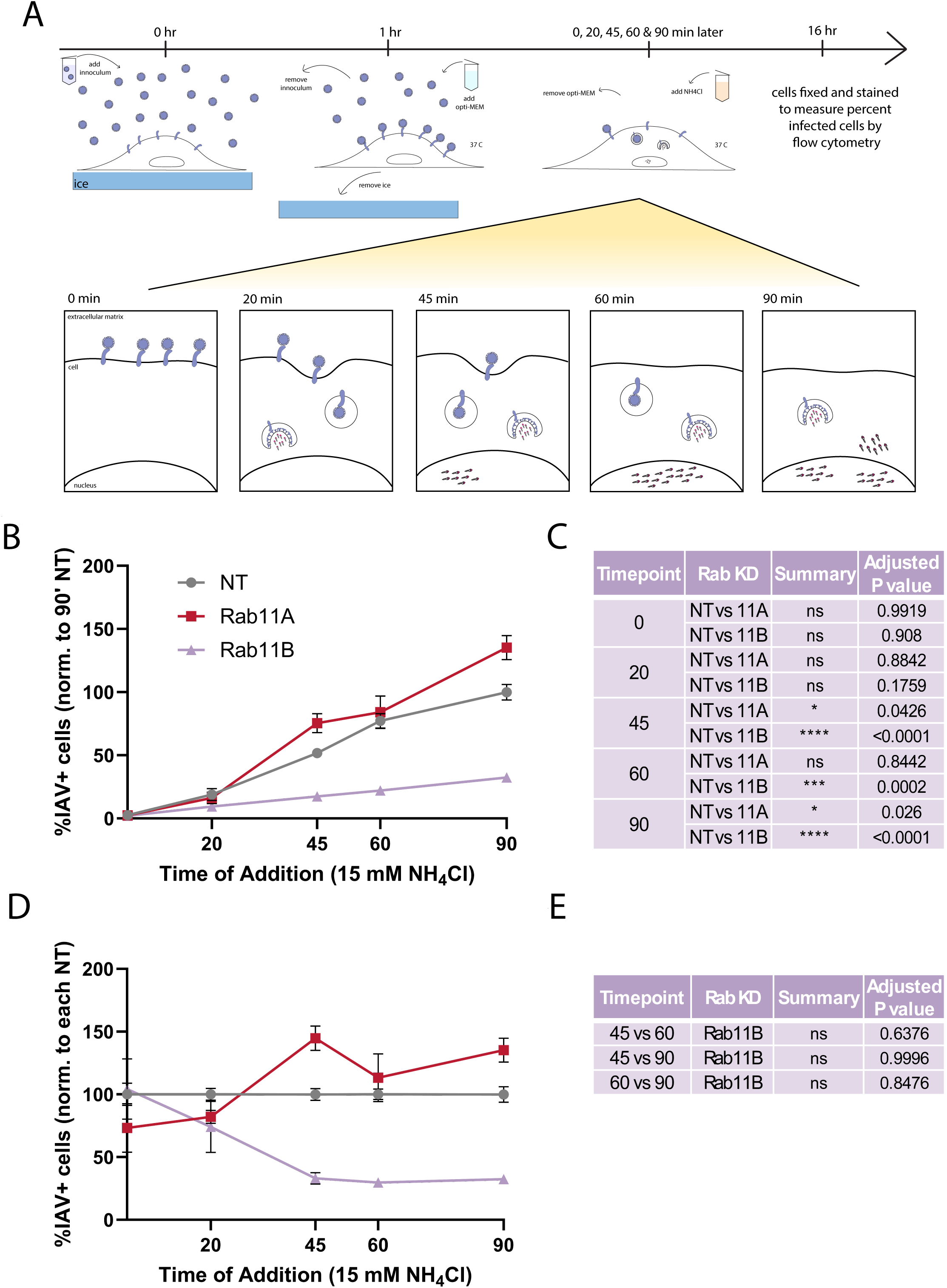
Loss of Rab11B delays the entry kinetics of the H3N2 IAV subtype. **A)** A549 cells were treated with siRNAs targeting Rab11A, Rab11B or a non-targeting control, 48 hpt cells were infected with 2022 IAV isolate of H3N2 (A/Burlington/UVM-1927/2022) at an MOI of 1, or mock infected. At the indicated times (0, 20, 45, 60 or 90 minutes) post infection, viral overlay was removed and replaced with 15 mM NH_4_Cl (to prevent endosomal acidification and thus fusion of internalized virions) for the remainder of the infection. 16 hours post inoculation cells were washed with PBS, trypsinized, fixed and stained for flow cytometry analysis to determine total (intracellular as well as extracellular) staining using anti-HA or anti-M2 antibodies. The percent of HA^+^/M2^+^ cells in each experiment was determined using the mock control to determine background staining levels. Mean +/- SEM is plotted, normalized to the average of the NT control for each biological experiment **B)** at 90 minutes or **D)** at each timepoint. **C)** Statistical comparisons of Rab11A or Rab11B depleted conditions vs the NT control were conducted using a two-way ANOVA with Dunnett’s multiple comparisons test and p values are reported. **E)** Statistical comparisons of the Rab11B depleted condition over time were conducted using a two-way ANOVA with Tukey’s multiple comparisons test and p values are reported. N=6 replicates from three biological experiments, [*(p<0.05), ***(p<0.001), ****(p<0.0001)].

Given the profound defects we observed in overall entry, we next sought to determine which viral gene (or genes) contributed to the singular dependence of our H3N2 viruses on Rab11B. Viral binding is governed primarily by HA (with a possible role for NA) with fusion dependent on HA, uncoating relying on M2, and transport to the nucleus likely mediated by multiple genes. After first verifying that A/Puerto Rico/8/1934 (H1N1, referred to as PR8 in future) was not dependent on Rab11B for viral protein production (Fig 7A-D), we created 7+1 reassortments of the UVM-1927 HA or NA gene segments in a PR8 background using an established reverse genetics system. The PR8 7+1 reassortant containing the UVM-1927 HA gene (PR8: HA_UVM-1927) showed a significant (75-90%) reduction in viral protein levels when HA, NP and M2 expression was quantified in cells lacking Rab11B (Figure 7E-H). Conversely, no significant reduction in viral protein expression was seen in 7+1 reassortant containing the UVM-1927 NA gene (Figure 7I-L). This data shows that the dependence on Rab11B for viral protein production can be mapped to the HA segment, and to the HA protein itself given that segment 4 is monocistronic^45^. This result strongly suggested that Rab11B is required at one of the earliest stages of entry, given that HA plays key roles in binding and fusion.

**Figure 7.**
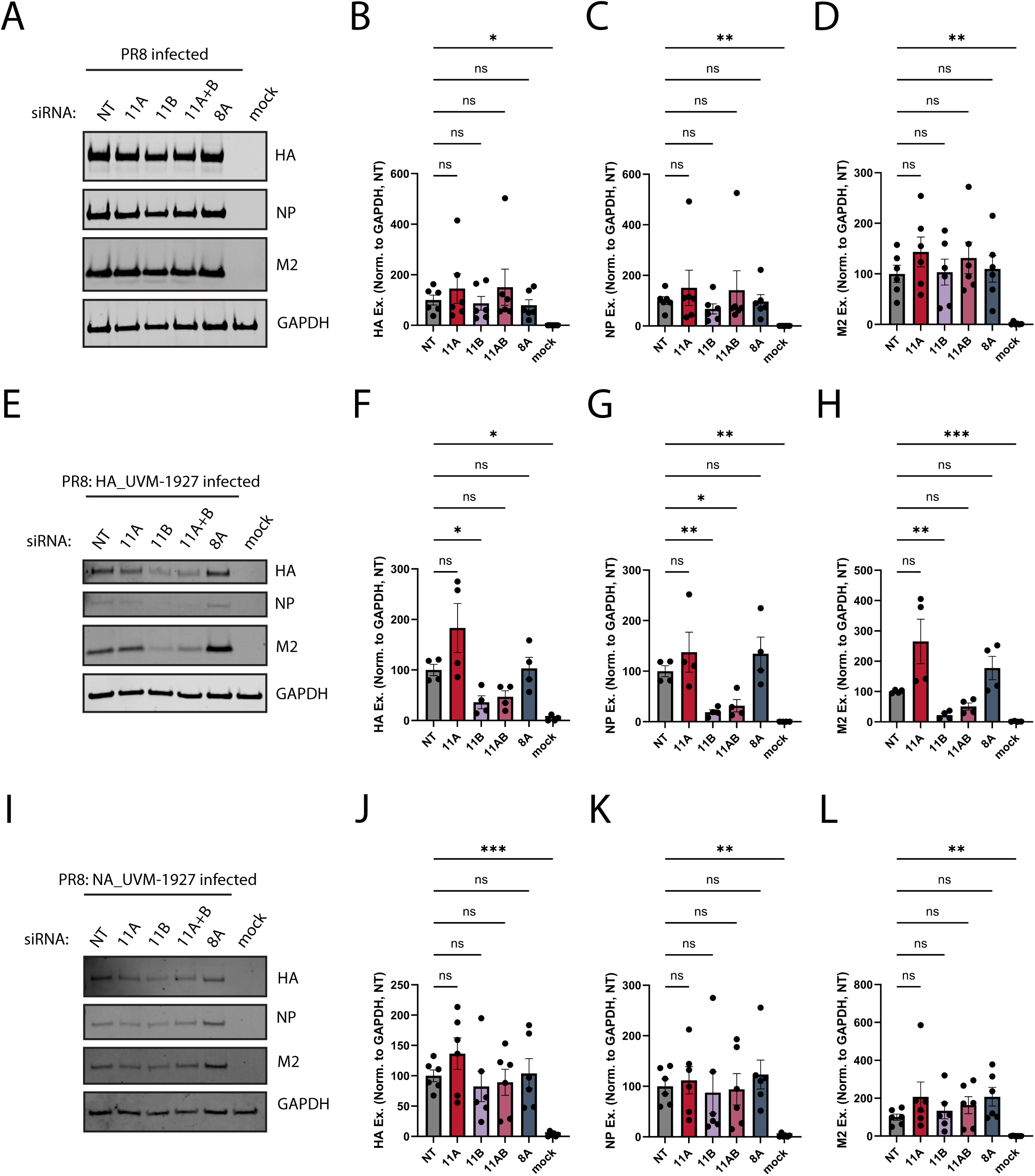
H3N2 dependence on Rab11B early in infection can be mapped to the HA gene. A549 cells were treated with siRNAs targeting Rab11A, Rab11B (singly and in combination), Rab8A or a non-targeting control, 48 hpt cells were infected with **A)** A/Puerto Rico/8/1934 (PR8), **E)** a reverse genetics derived 7+1 reassortment containing the HA gene of UVM-1927 (PR8:HA UVM-1927) or **I)** a reverse genetics derived 7+1 reassortment containing the NA gene of UVM-1927 (PR8:NA UVM-1927) at an MOI of 1, or mock infected. 16 hpi, cell lysates were collected and visualized by SDS-PAGE and western blot using rabbit anti-HA and anti-GAPDH antibodies in addition to mouse anti-NP and anti-M2 antibodies. Expression of viral proteins was quantified and normalized to GAPDH levels and the average of the NT controls for each biological replicate is shown for PR8 (**B, C, D**), PR8:HA UVM-1927 (**F, G, H**) and PR8:NA UVM-1927 (**J, K, L**). (HA0, **D)** NP and **E)** M2. Mean +/- SEM is plotted, [*(p<0.05), **(p<0.01), ***(p<0.001), ****(p<0.0001)], N=6 from three biological experiments (B, C, D, J, K, L) or 4 from two biological experiments (F, G, H). Statistical comparisons done using Welch’s one way ANOVA with Dunnett’s multiple comparisons.

To explore our hypothesis that Rab11B was acting at the earliest stages of entry, we used an RT-qPCR based assay to examine how the rate of binding was affected by depletion of Rab11B. We adapted an assay that we have previously used to measure levels of Hantavirus binding^46^, in which we synchronized viral binding to the surface of cells by binding at 4°C to prevent endocytosis. We then washed cells to remove unbound virions and used RT-qPCR to measure the viral genome copies of UVM-1927 (H3N2) attached to cells after one hour. To verify that we were measuring authentic bound virus (rather than virus sticking non-specifically to cells or plastic), we used exogenous neuraminidase (NA) to cleave bound virus, as NA plays a key role in cleaving sialic acids to release newly produced virions that would otherwise remain attached to the producer cell^21^. Using this control, we found a significant drop in viral binding, with ∼97% of viral RNA removed upon NA treatment (Figure 8A). Having confirmed we were detecting authentically bound virions, we next conducted the binding experiment with UVM-1927 in cells lacking Rab11A or Rab11B. Notably, in cells lacking Rab11B we observed a significant (∼50%) decrease in binding compared to cells treated with a NT control, while binding of virions in cells depleted of Rab11A was unaffected (Figure 8B). Next, we repeated this experiment using the UVM-0478 (H1N1) virus, to confirm that the binding defect was specific to the scenario in which viral protein production failed to occur. In this scenario, cells lacking Rab11B had no defect in viral binding (Figure 8C), suggesting that the ability of Rab11B to support viral binding is specific to H3N2 viruses, and is dispensable for the binding of this H1N1 isolate.

**Figure 8.**
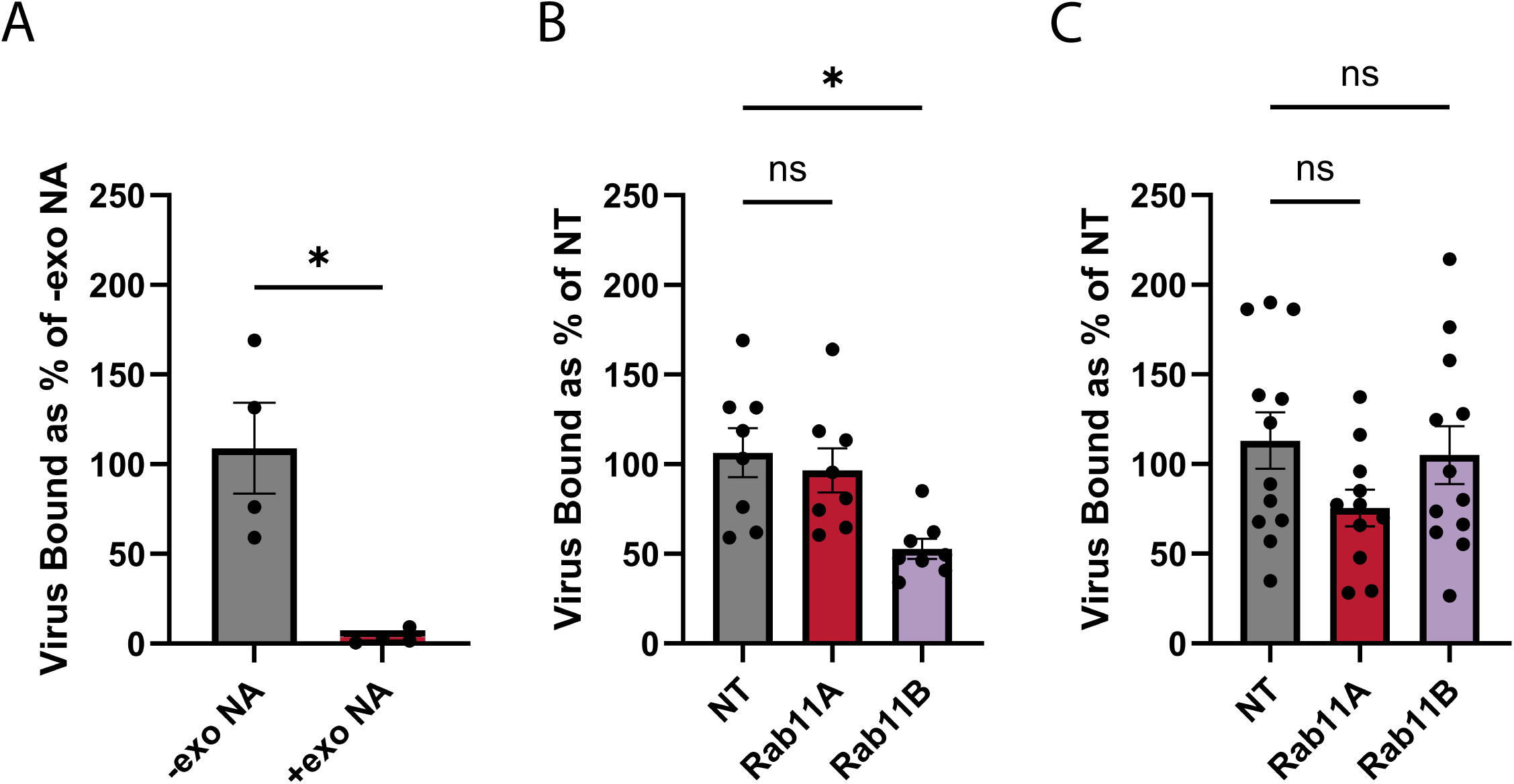
Rab11B is required for efficient binding of H3N2 but not H1N1 virions to the cell surface. A549 cells were prechilled on ice after which cold virus (A/Burlington/UVM-1927/2022 [H3N2] at an MOI of 1) was bound to cells for one hour to synchronize binding. Cells were washed three times with ice cold PBS to remove unbound virus, and total virion binding was determined by extracting RNA from each well and using RT-qPCR to determine relative levels of IAV RNA vs a housekeeping control (18S). **A)** Bound virions were stripped by treating cells with exogenous neuraminidase. Alternatively, cells were treated with siRNAs targeting Rab11A, Rab11B or a non-targeting control for 48 h before binding **B)** a H3N2 (A/Burlington/UVM-1927/2022) or a **C)** A/Burlington/UVM-0478/2022 as above. Mean +/- SEM is plotted, normalized to the average of the NT control in each biological experiment [*(p<0.05)], N=4 from two biological experiments **(A)**, N=8 from three biological experiments **(B),** N=11-12 from two biological experiments **(C).** Statistical comparisons done using the Welch’s t Test (A) and the Brown-Forsythe and Welch ANOVA tests.

## DISCUSSION

The data presented here are consistent with a model in which recently circulating H1N1 and H3N2 influenza subtypes bind to different cellular receptors or attachment factors. In this framework, the H3N2 receptor(s) is trafficked by Rab11B, while the H1N1 receptor(s) is transported by a non-Rab11A/B dependent mechanism (Figure 9). While we were initially surprised to observe such different roles for these two Rab11 isoforms in the context of influenza infection, prior literature does support the isoforms playing distinct and even opposing functions within the cell, despite their sequence similarity^42,43^. To some degree, the distinct functions of Rab11A and Rab11B are a result of differential expression, where Rab11A is ubiquitously expressed while Rab11B is enriched in the brain, testis and heart (while having a lower but broad level of expression in other tissues)^43^. And in many cases, Rab11A and Rab11B can transport the same cargo, functioning in a redundant manner^47^.

**Figure 9.**
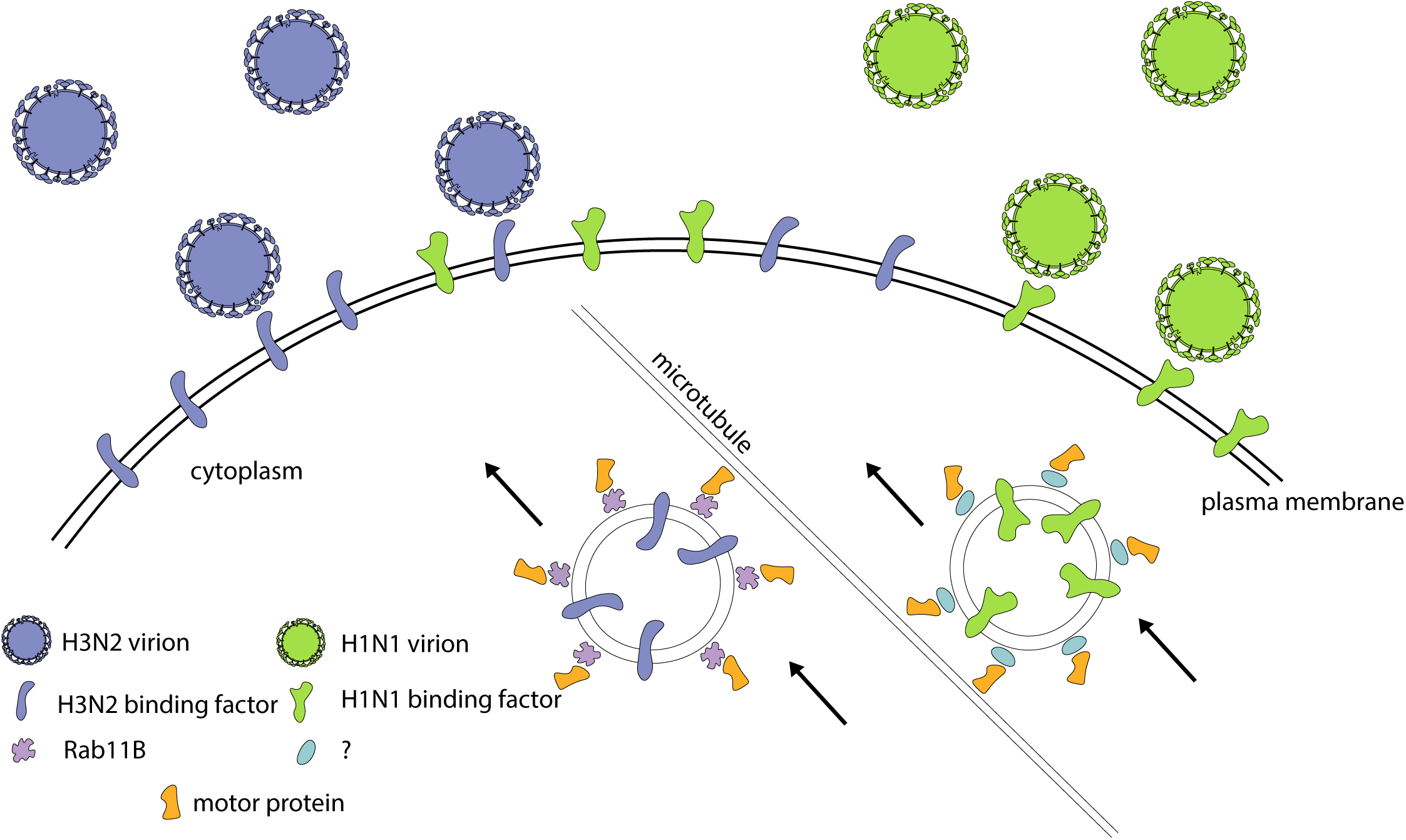
Rab11B may transport a H3N2 specific IAV binding receptor. We propose a model in which H3N2 virions (blue) bind to different cell surface proteins than H1N1 virions (green), and these cell surface proteins (putative binding ‘receptors’) are differentially transported by the Rab11B recycling pathway. In this framework, protein(s) required for H3N2 virions to bind are transported on Rab11B-positive vesicles, while the molecule(s) required for H1N1 binding are transported by a different (as of yet unidentified) cellular pathway that is independent of Rab11B.

However, there are also examples of Rab11A and Rab11B playing distinct roles within the same cellular background. Rab11B plays a role in trafficking fibroblast growth factor receptor 4 (FGFR4) that is different and distinct from Rab11A’s role in this pathway^48^, and Rab11B but not A have been implicated in transport of the cystic fibrosis transmembrane conductance regulator (CFTR)^49^. Notably Rab11B (but not Rab11A) is critical for recycling the Protease-activated receptor-1 (PAR1) while enhanced degradation of PAR1 observed in cells lacking Rab11B is blocked by simultaneous depletion of Rab11A^43^. This phenotype is similar to what we observed in our initial infection of cells with UVM-1927 (our H3N2 subtype), where depletion of Rab11B alone blocked viral protein production but simultaneous depletion of Rab11A + Rab11B rescued the phenotype (Figure 2E, S3). While Rab11A and Rab11B share a high degree of sequence similarity, the crystal structures of the two isoforms reveal potentially significant differences, as Rab11A is seen as a dimer in the crystal structure, while Rab11B crystallized as a monomer^50^. In addition, there were significant differences in the confirmation of the Switch I region, which is of particular interest given the likely role Rab11A’s Switch I region plays in binding to PB2^30,50^.

While it seems clear that Rab11A and Rab11B can transport distinct pools of proteins to the plasma membrane, the fact that Rab11B traffics a protein (or proteins) that is required for binding of H3N2 but not H1N1 subtypes of influenza virions is particularly interesting given our current understanding of influenza entry. Conventional thinking of influenza entry typically posits that influenza virions ‘roll’ along the cellular surface, interacting with sialoglycans present on various glycoproteins and glycolipids which mediates the first step, ‘attachment’. After this binding event, more recent work proposes that glycan-binding is followed by secondary engagement of the receptor, followed by internalization^18^. Within this framework, it has long been recognized that the linkage type and subsequent ‘shape’ of sialic acid modifications play important roles in the ability of a given HA to bind to a particular host cell, with α2,6 linked structures forming an ‘umbrella’ like structure preferred by human IAVs while α2,3 linkages form a ‘cone’ like shape that is preferentially recognized by strains of avian origin^51–58^.

Importantly, further modifications to the glycan also affect glycan topology, and it is known that not every glycan with a α2,6 linkage is recognized by every influenza strain that possess α2,6 receptor specificity^18^. While some studies report relatively similar glycan usage between seasonal H1N1 and H3N2 isolates, H3N2 isolates often bind a smaller subset of glycans^58,59^. H3N2 isolates are known to be particularly sensitive to passage in eggs, with H3N2 isolates developing increased specificity for α2,6 linkages, which has posed a problem for egg-grown vaccine development^60^. In addition, there are non-sialic acid attachment factors that can affect HA binding, such as phosphorylated but non-sialylated glycans^61^. The H17N10 and H18N11 influenza subtypes (found in South and Central American bats) cannot bind sialic acid^62–64^ and instead used the MHC-II receptor^65^. Furthermore, H2N2 viruses from both human and avian origin have dual receptor specificity and are able to utilize either a sialic-acid dependent or a sialic acid-independent, but MHC-II dependent, mechanism of entry^66^.

Some studies have also shown that specific cell surface proteins have the ability to interact with HA in a sialic acid-independent manner, including nucleolin^67^ and natural killer (NK) cell p46-related protein (NKP46)^68–70^. Other work concludes that a specific sialyated protein is responsible for HA binding and subsequent internalization, with the voltage-dependent CA^2+^ channel Ca_v_1.2 reported to play this role for PR8^8^. Engagement of cellular signaling receptors, including phosphatidylinositol-3-kinase and receptor tyrosine kinases (RTKs) including the epidermal growth factor receptor (EGFR) have also been shown to play a role in transmitting signals across the plasma membrane that lead to internalization^9–14^. Finally, there are reports of subtype specific dependence on cellular entry factors, with post-binding internalization of H1N1 but not H3N2 subtypes depending on the phospholipase C-γ1 (PLC-γ1) signaling mediator downstream of receptor tyrosine kinase pathways^15^. As much prior work has focused on binding and entry of H1N1 strains (typically using the highly lab adapted PR8 strain), it is likely that further subtype specificities in cellular binding and internalization receptors remain to be identified.

Our data clearly establishes the subtype specific role of Rab11B in the binding of recent seasonal H3N2 but not H1N1 isolates. However, as Rab11B is not a transmembrane protein, it almost certainly plays an indirect role in this process by trafficking a protein or proteins required for H3 binding to the cell surface. Given the abundance of proteins in the surface proteome with an α2,6 linkage, it seems somewhat unlikely that Rab11B is solely responsible for trafficking the entire variety of proteins conventionally thought to serve as the binding receptor for human IAV strains. Further arguing against a scenario in which the loss of Rab11B is simply reducing cell surface levels of sialydated proteins is the fact that entry and binding of H1N1 isolates from the same time period are unaffected by the Rab11B depletion. Our data therefore supports the existence of a more specific protein-based binding-receptor that is differentially used by H3N2 isolates from recent years (Figure 9). Open questions that remain the subject of future study include determining whether the Rab11B-dependent cargo that is transported to the surface is responsible for mediating internalization of the virion in addition to binding. Mapping the plasma membrane resident cargos of the Rab11B-dependent recycling pathway could provide a valuable starting point in identifying putative attachment and/or internalization receptors used specifically by recent H3N2 IAVs. Future work is also needed to determine whether these findings extend to older H3N2 isolates, as well as whether this observation is true in cell types of other origins and species. In summary, this work extends the well-established dependence of IAV on Rab11a late in infection to include recent seasonal H1N1 and H3N2 isolates, while also extending our understanding of the critical role this protein family plays to include Rab11b mediated binding and entry of H3N2 viruses.

## MATERIALS AND METHODS

### Ethics Statement and Institutional Approvals

Research conducted in this study was reviewed and approved by the Institutional Biosafety Committee of the University of Vermont (REG202100008). The use of deidentified positive clinical specimens was approved by the University of Vermont Institutional Review Board (STUDY00000881) under a waiver of consent.

### Cells

Human adenocarcinoma 549 cells (A549) (kindly obtained from Dr. Steve Baker), human embryonic kidney cells (HEK-293T/17) (kindly provided by Dr. John Salogiannis), human papillary adenocarcinoma lung cells, considered ‘club-like cell’ (NCI-H441, ATCC HTB-174) and Madeline Darby Kidney cells modified to express α2,6 sialic acid and TMPRSS2^71^ (MDCK-SIATT) (kindly provided by Dr. Jesse Bloom via Dr. Steve Baker) were cultured in 1X DMEM (Corning #06923002) with 10% FBS (Gibco #16140-071) and 1X pen/strep (Corning #30-002-CI). NCI-H441 cells stably transduced with Rab11B targeting shRNA were maintained in media as above, plus 3 μg/mL puromycin.

### Viruses

A/Burlington/UVM-1927/2022 (H3N2) (UVM-1927) and A/Burlington/UVM-0478/2022 (H1N1) (UVM-0478) viruses were isolated from IAV positive clinical specimens graciously provided by Dr. Jessica Crothers at the University of Vermont. MDCK-SIATT cells were seeded in a 12-well plate (CytoOne #CC7682-7512) at a density of 3.5x10^5^ cells per well 24 hrs before isolation. Day of isolation, each clinical sample was brought up to a total volume of 400ul in serum-free media (Corning #10-017-CM). Plated cells were washed twice in PBS (Corning #21-040-CV) and 400ul of sample placed in each well. Cells were incubated for 1hr at 37°C and 5% CO_2_, rocking the plates every 20 mins during the incubation. After 1hr, the supernatants were removed from each well and replaced with 1mL media comprised of Opti-MEM (Gibco #11-058-021) + 1ug/mL TPCK-Trypsin (Sigma-Aldrich #T88002-100MG) + 1X pen/strep (Corning #30-002-CI). Samples were deemed ready to be collected when visible cytopathic effect (CPE) could be seen. At that time, supernatants were clarified at 10,000G for 5 minutes, transferred to clean tubes and stored at -80°C for later stock generation. Low passage stock of A/Baltimore/JH-0586/2022 (H3N2) (JH-0586) virus (EPI_ISL_16766241) originally isolated in human nasal epithelial cells was a kind gift of Dr. Andrew Pekosz at Johns Hopkins University. A/Puerto Rico/8/1934 (H1N1) was derived by reverse genetics.

### Generation of 7+1 Reassortants by Reverse Genetics

Viruses containing the HA or NA segments of A/Burlington/UVM-1927/2022 (H3N2) in a A/Puerto Rico/8/1934 (H1N1) backbone were created by reverse genetics, using a pHW2000 plasmid system kindly provided by Dr. Stacey Schultz-Cherry. The HA or NA coding sequence (based on sequencing of the UVM-1927 stock) was synthesized and cloned into the pHW2000 HA or NA segment plasmid (replacing the original PR8 coding sequence) by Genscript. HEK-293T cells were seeded in six well plates (7x10^5^ cells/well) in complete DMEM. The following day cells were transfected with 450ng of each of the eight pHW2000 plasmids (PB1, PB2, PA, HA, NP, NA, M, NS) using jetOPTIMUS (for per reaction volumes of 3.5 ug combined segment DNA, 50 ul jetOPTIMUS buffer, 3.5 ul jetOPTIMUS, incubated for ten minutes before dropwise addition to adherent HEK-293T cells). 24 hours post transfection media was removed and HEK 293T cells were overlaid with 3x10^5^ MDCK-SIATT cells/well resuspended in 2 ml OPTI-VGM (optiMEM, 0.19% BSA, 1X Pen/Strep, 1 ug/ml TPCK Trypsin). Cells were monitored every 24 hr for cytopathic effect compared to a transfection control lacking PB2. Once CPE was observed, rescued virus was blind passaged by infecting a confluent T25 of MDCK-SIATT cells with 500 ul clarified rescue supernatant combined with 500 ul OPTI-VGM, 2 ug/ml TPCK-trypsin for three days. Blind passages were titered by plaque assay and used to generate additional high titer stocks.

### Viral Stocks

Viral stocks were grown by seeding 4.2 x 10^6^ cells/flask of MDCK-SIATTs in a T150 flask (Corning #430825) in 1X DMEM (Corning #06923002) with 10% FBS (Gibco #16140-071) and 1X pen/strep (Corning #30-002-CI) and incubated overnight at 37°C and 5% CO_2_. Each flask was washed twice with PBS (Corning #21-040-CV) to remove serum and infected at an MOI of 0.001 in a volume of 5 ml for one hour, overlaid with 15 ml of Opti-MEM (Gibco #11058-021) plus 1 ug/ml TPCK trypsin (Sigma-Aldrich #T88002-100MG) and incubated at 37°C until ∼50% CPE was observed.

### Viral Infections

Viral infections were performed at the indicated MOI, by first washing the cells once with PBS (Corning #21-040-CV) and then infecting with an inoculum made up of virus plus 1X DMEM (Corning #06923002) at 37°C and 5% CO_2_ for one hour. Cells were rocked every 20 minutes, and after one hour the inoculum was removed and replaced with an overlay of Opti-MEM (Gibco #11058-021) before incubating for the indicated time at 37°C. Viral supernatants were clarified by centrifuging at 10,000G for 5 minutes to remove any cell debris, aliquoted into two separate tubes and stored at -80°C.

### Viral Plaque Assays

Infectious viral production was assessed using plaque assays. MDCK-SIATT cells were trypsinized using 0.25% trypsin-EDTA (Gibco #25200-0720) for 10-15 minutes and seeded at 1x10^6^ cells/well of a six well plate (Corning #3506) The following day wells were washed with PBS (Corning #21-040-CV) or DMEM (Corning #06923002) and inoculated with serial ten-fold dilutions of each sample in DMEM. After infection, cells were placed at 37°C with 5% CO_2_ for an hour and rocked every 15 minutes to ensure even mixing and prevent cell death from drying. After one hour, cells were overlayed with 2 ml of a warmed 1:1 mixture of 2.4% Avicel RC-591 NF (Dupont #RC591-NFDR080) + 1X DMEM (Corning #06923002) + 1 ug/ml TPCK trypsin (Sigma-Aldrich #T88002-100MG). Cells were returned to 37°C for 48 hours, and care was taken not to disturb the plates during this period. Finally, cells were washed with PBS (Corning #21-040-CV), fixed with 4% formaldehyde (Honeywell #F16354L) for 20 minutes and stained with 0.1% crystal violet (Fisher #C581-100) for five minutes. Plates were rinsed three times with water and allowed to dry before plaques were counted to determine viral titer.

### Viral Sequencing

Viral subtypes were identified through amplicon based whole genome sequencing. In brief, RNA from viral stocks was extracted using a QIAamp Viral RNA Mini Kit (Qiagen #52906) according to manufacturer’s protocols, and one step RT-PCR was used to amplify all eight segments using universal primers^72^ and Superscript III high-fidelity RT-PCR kit (Invitrogen #12574-035).

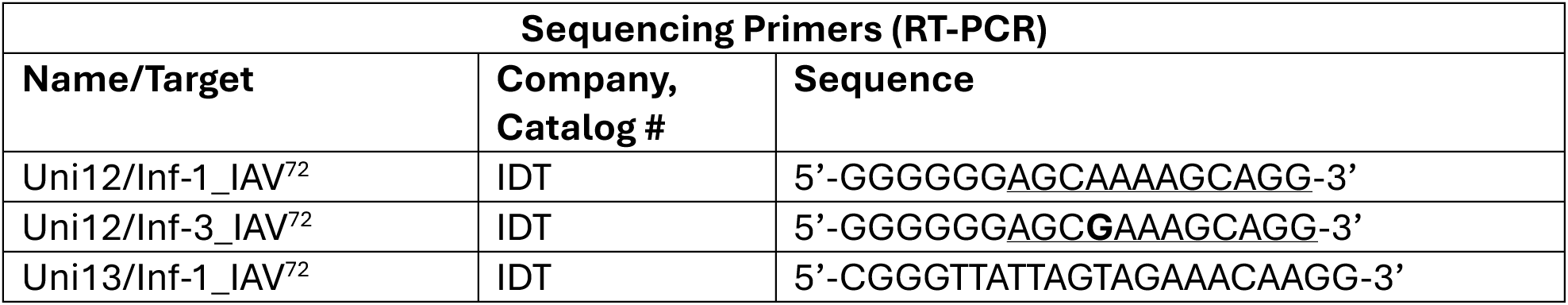

Per reaction, 8.75 ul nuclease free water, 12.5 ul of 2X RT-PCR buffer 0.2 ul of 10 uM Uni12/Inf-1, 0.3 ul of 10 uM Uni12/Inf-3, 0.5 uL of 10 uM Uni13/Inf-1, and 0.25 ul RT/Taq enzyme mix were combined with 2.5 ul of RNA template. RT-PCR was run on a Eppendorf Mastercycler Nexus Thermal Cycler with the following cycling conditions: 55°C/2 min, 45°C/60 min, 94°C/2 min, 5 cycles of [94°C/30s; 44°C/30s; 68°C/3.5 min], 26 cycles of [94°C/30s; 57°C/30s; 68°C/3.5 min], 68°C/10 min, hold at 4°C.

Successful amplification was confirmed by visualizing PCR products (i.e., seven-eight bands) on a 1% agarose (Fisher #BP160-500) gel for 2 hours at 120V. Samples were then purified using a QIAquick PCR Purification Kit (Qiagen #28104) and Illumina based amplicon sequencing conducted by the Emory EPC Genomics Core.

We used a Nextflow pipeline developed by Dr. Ramiro Barrantes Reynolds at the Vermont Integrative Genomics Resource Core. This pipeline uses Nextflow/NF-Core based UPHL-BioNGS/Walkercreek v2.0.0 bioinformatics pipeline for analysis and adds a custom Nextflow pipeline for generating a phylogenetic tree^73–79^. We compared the similarity between our two H3N2s by using an RShiny app developed to examine the results of the pipeline described above and understand the sequence level differences between the two viruses. We identified their relatedness through the tree created by the pipeline described above^80^. All R coding was done in RStudio^81^ (R2023.09.1+494) and we used Nextflow 24.04.4 for pipeline development. Computation was done using the Vermont Advanced Computing Center (VACC).

### siRNA knockdowns

A549 cells were seeded in 24 well plates at a density of 1.5x10^5^ cells/well, in a volume of 500 ul. Transfections were performed on cells in suspension or within 24 hours of plating. To knockdown our gene of interest, we transiently transfected siRNA sequence targeting Rab11A, Rab11B, Rab11A+B, Rab8A or a non-targeting siRNA, using Lipofectamine RNAiMAX Reagent (Invitrogen #56532) in A549 cells. For each well, we combined 1.2 uL of 10 uM siRNA stock with 198.6 ul of Opti-MEM (Gibco #11058-021) and 2 ul of RNAiMax (Invitrogen #56532), gently mixed and allowed to rest for 20 minutes before adding to cells dropwise. 48 hours post transfection cells were either infected, or RNA was harvested to verify mRNA knockdown at the time of infection.

| siRNA Target | Company, Catalog # | Assay ID |
| --- | --- | --- |
| Non-targeting control | ThermoFisher, 4390843 | NA |
| Rab11A | ThermoFisher, 4390824 | s16704 |
| Rab11B | ThermoFisher, 4390824 | s17648 |
| Rab8A | ThermoFisher, 4390824 | s8681 |

### shRNA depletion

#### Lentivirus Generation

A custom lentiviral construct driving inducible shRNA expression targeting human Rab11B was obtained from Horizon Discovery (SMARTvector inducible human Rab11B mCMV-Turbo RFP shRNA). Lentiviral particles were generated by transfecting subconfluent HEK-293T cells with 6 ug of the Rab11B SMARTvector construct and 28.5 ug of the Horizon Trans-Lentiviral shRNA packaging kit (Horizon Discovery, cat # TLP5912) using jetOPTIMUS (Sartorius Item # 101000025) according to the manufacturer’s instructions. Cells were incubated for 16 hrs before transfection media was changed to reduced serum media (1X DMEM containing 5% FBS, 1X Pen/Strep). For each of the following three days, supernatant was collected and replaced with fresh media. The supernatants were clarified and stored at -80°C for use in subsequent transductions.

#### Lentiviral transduction

H441 cells were seeded at 1.5x10^5^ cells/well in a 24 well plate and transduced the following day with lentiviral supernatants by spinnoculation. H441 cells were inoculated with 500 ul total volume (consisting of complete DMEM, 0.5 μl polybrene, and 100μl of lentiviral supernatant). Cells were spun for two hours at 640xG, 20°C, after which cells were returned to the incubator. The following day, media was replaced, and cells were returned to the incubator. Three days post transduction cells were selected with puromycin (3 μg/mL), the dilution in which ∼30% of cells survived was maintained in puromycin containing media and expanded for future use.

#### Silencing and validation

Rab11B depletion was optimized by comparing a range of doxycycline concentrations (0.1 μg, 0.5 μg, 2 μg, 4 μg) and times post induction (24, 48, 72 hrs). Rab11B silencing was measured by RT-qPCR, cells were lysed in 350 μl of RLT buffer for RNA extraction and RT-qPCR as described below. The optimal time and concentration was found to be 4 ug/mL of doxycycline 72 hrs post induction so this was used for all further shRNA depletion experiments.

### RNA extraction

48 hrs after transfection, cells treated with siRNAs were lysed in 350 ul of RLT lysis buffer (from RNeasy Mini Kit). Cell lysates were processed, and RNA was extracted using the QIAshredder columns (Qiagen #79656) followed by RNeasy Mini Kit (Qiagen #74106) coupled with on column DNase treatment using RNase-Free DNase Set (Qiagen #79256), according to manufacturer’s protocols.

### RT-qPCR

We used a 20 ul reaction consisting of 5 ul of New England Biolabs Luna Probe One-Step RT-qPCR 4X Mix with UDG (NEB #M3019E), 1 ul of the 20X primer/probes stock, 1 ul of extracted RNA, and 13 ul of nuclease-free water for all of our RT-qPCR with pre-mixed primer/probes (listed below). To measure levels of IAV, we used the SVIP-MPv2 primers and probe^82^. We again used 20 ul reaction volume with 5 ul of New England Biolabs Luna Probe One-Step RT-qPCR 4X Mix with UDG (NEB #M3019E), 0.28 ul each of 100 uM forward and reverse primer stock, 0.08 ul of 100 uM probe stock, 13.36 ul of nuclease-free water, and 1 ul of template RNA. Each sample was run in triplicate, and each primer/probe set had a no template control. Once loaded into plates (Applied Biosystems #4306737), samples were spun at 400 g for 5 minutes. RT-qPCR was run using a QuantStudio Real-Time PCR system using the following cycling conditions: 55°C/10 min, 95°C/1 min, 45 cycles of [95°C/10s; 60°C/30s], 4°C hold. Relative RNA levels were determined using the ΔΔC_T_ method, using 18S RNA levels as the housekeeping control.

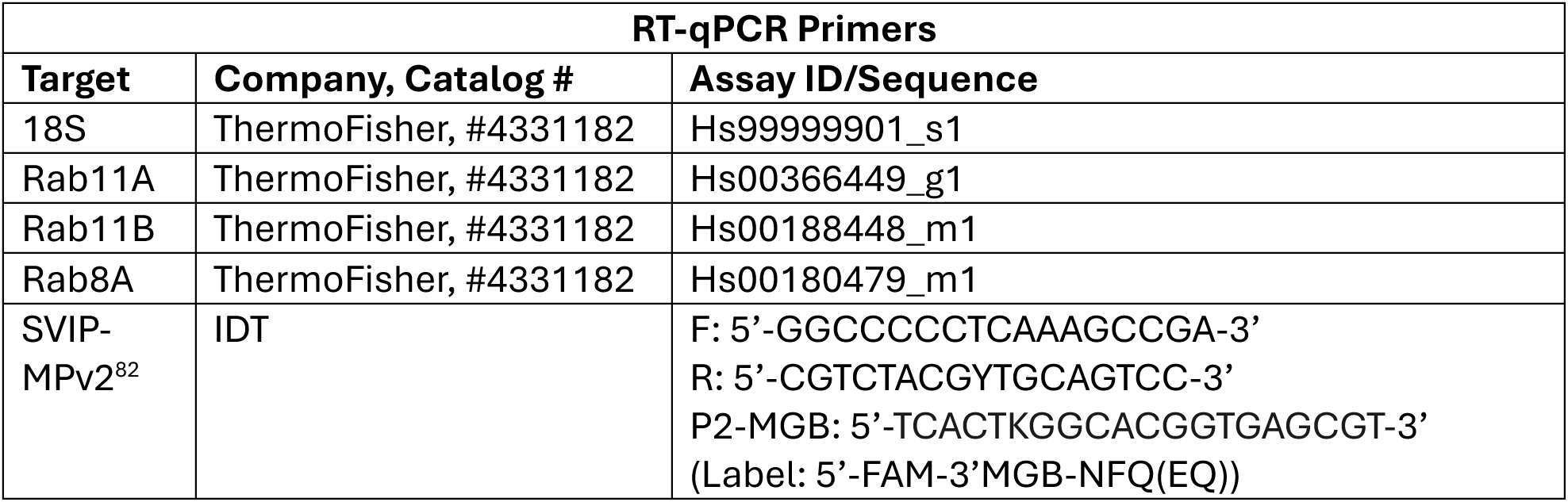

### Antibodies

Antibodies.
| Primary Antibodies |  |  |
| --- | --- | --- |
| Species, Target | Company, Catalog # | Dilution Used |
| Rabbit anti-HA | Sino Biological<br>#86001-RM01 | 1:5,000 (western blot)<br>1:1,000 (flow cytometry) |
| Mouse anti-M2 (14C2) | Abcam #AB5416-1001 | 1:5,000 (western blot)<br>1:1,000 (flow cytometry) |
| Mouse anti-NP (HB65) | BioXCell<br>#BE0159R001MG | 1:1000 (western blot) |
| Anti-GAPDH | Abcam #AB181602 | 1:10,000 (western blot) |
| Rabbit anti-Udorn (H3N2) | From Dr. Robert Lamb | 1:1,000 (flow cytometry) |
| Secondary Antibodies |  |  |
| IRDye 680LT goat anti-rabbit IgG | LICOR #92668021 | 1:10,000 |
| IRDye 680RD goat anti-mouse IgG | LICOR #92668070 | 1:10,000 |
| IRDye 800CW goat anti-mouse IgG | LICOR #92632210 | 1:10,000 |
| Alexa Fluor 488 anti-rabbit | Invitrogen #A11034 | 1:1,000 |
| Alexa Fluor 647 anti-mouse | Invitrogen #2555690 | 1:1,000 |

### Protein analysis

Samples were prepared for protein analysis and western blot largely as previously published (Kubinski, 2024). Briefly, cells were lysed in a NP-40 (Alfa Asear #J60766) + 1% Triton X-100 (Fisher #BP151-100) supplemented with Pierce Protease Inhibitor Mini Tablets (Thermo Scientific #A32955) for 20 mins on ice. Cell lysates were clarified, then diluted 1:1 (v/v) with 4X Laemmli sample buffer (250 mM Tris-HCl pH 6.8, 40% glycerol, 8% SDS, 0.04% Bromophenol Blue, 2.75mM 2-mercaptoethanol). All samples were heated to 95°C for 5-10 mins before loading. Samples were separated in NuPAGE 4–12% Bis-Tris gels (Invitrogen # NPO335BOX) in MES buffer (Invitrogen # NP0002) with a molecular mass ladder (Thermo Fisher #LC5925) at 180v for 50 mins before being transferred into a nitrocellulose membrane (Invitrogen # IB23001) using an iBlot 2 machine (Invitrogen) at 20V for seven minutes.

Membranes were blocked in a 5% milk/PBS solution for 30 minutes and then incubated overnight at 4°C in a solution containing the primary antibody in a 5% milk/PBST (PBS + 0.2% Tween 20). Membranes were washed three by five minutes in PBST, incubated while rocking with secondary antibodies diluted in 5% milk/PBST for 45 minutes, washed again with PBST and imaged with a LI-COR Odyssey CLx. Protein expression was analyzed by measuring band densitometry in the LI-COR software package, Image Studio (Ver 5.5).

### Flow Cytometry

Infected cells were washed in PBS, trypsinized, and fixed in a final concentration of 4% paraformaldehyde (Thermo Scientific #043368.9M) for 20 minutes. After fixing, cells were transferred to 5 mL polystyrene round-bottom tubes (Falcon #352052) and spun at 400g for 5 minutes to pellet cells. Supernatants were discarded and cells resuspended in 4 mL of wash buffer (1X PBS/5% FBS). Cells were pelleted and 200 ul of PBS/0.05% Triton X-100 (Fisher #BP151-100) solution was added to permeabilize for five minutes. Cells were pelleted and washed as above before resuspending in 1 mL of wash buffer for one hour to block. Cells were pelleted and resuspended in 200 ul of primary antibody solution (diluted in wash buffer) for one hour. Cells were pelleted and washed as above before resuspension in 200 ul of secondary antibody solution (again diluted in wash buffer) for one hour (protected from light). Cells were washed and pelleted one final time after which they were resuspended in 200 ul of sterile PBS and stored at 4°C protected from light until flow cytometry was performed. Samples were run on a Beckman CytoFlex (2L/4 flourescences) in the Harry Hood Bassett Flow Cytometry and Small Particles Detection facility (RRID:SCR_022147) at the UVM-LCOM and analyzed using FloJo software (v10.10.0).

### Binding Assay

Cells that had been transfected with siRNA 48 hrs earlier were placed on ice to cool, washed with ice cold PBS and infected with virus at an MOI of 1 (diluted in a binding buffer of Opti-MEM, 10 mM HEPES,10 mM MES to prevent changes in pH during incubation without CO_2_). Cells remained on ice (to prevent endocytosis), virus was bound for one hour and rocked every 15 mins to ensure cells an equal distribution of viral inoculum and prevent cells from drying. Cells were then washed three times with ice cold PBS to remove unbound virions. At this point, cells were either lysed in 350 ul RLT buffer (for RNA extraction and analysis of bound virions) or used for the time of addition assay. Alternatively, cells were treated with 0.0043 U/ul of exogenous neuraminidase (Roche #11585886001) for 20 minutes at 37°C to strip bound virions from the cell surface^21^. Cells were washed three times with PBS after neuraminidase treatment before lysis in 350 ul RLT as above.

### Time of Addition

Using a protocol adapted from one recently described by the Chlanda lab ^44^, we first synchronized infection using the binding protocol described above. After the initial hour of viral binding, cells were overlaid with Opti-MEM and were placed at 37°C. At 0, 20, 45, 60, or 90 mins, the Opti-MEM was removed and replaced with Opti-MEM containing 15 mM ammonium chloride (NH_4_Cl). Cells were then incubated at 37°C for 16 hrs, before preparing for flow cytometry as described above.

## Acknowledgements

We would like to thank Drs Edward Hutchinson, Margaret Kielian, Chris Huston, John Salogiannis, and Bruno Martorelli Di Genova for helpful discussion, Dr. Stacey Schultz-Cherry for the PR8 reverse genetics system, and Dr Andrew Pekosz for providing the A/Baltimore/JH-0586 (H3N2) viral isolate. All flow cytometric (small particle detection) data were carried out in the Harry Hood Bassett Flow Cytometry and Small Particles Detection facility (RRID:SCR_022147) at the UVM-LCOM and we thank Dr. Roxana del Rio-Guerra for her technical assistance with flow cytometry. The authors acknowledge the Vermont Advanced Computing Center (VACC) at the University of Vermont for providing computational resources that have contributed to the research results reported within this paper.

## Funding

This study was supported by the Microbiology and Molecular Genetics Department 2024 Distinguished Undergraduate Research Summer Award (A.H.T.), NIH K23AI175660 (J.W.C.), NIH (NIGMS) award P20GM125498 (E.A.B) and UVM start-up funds (E.A.B.).

## Data Availability

Raw sequencing data are available in the NCBI Bioproject ID PRJNA1254704.

---

**Figure S1.**
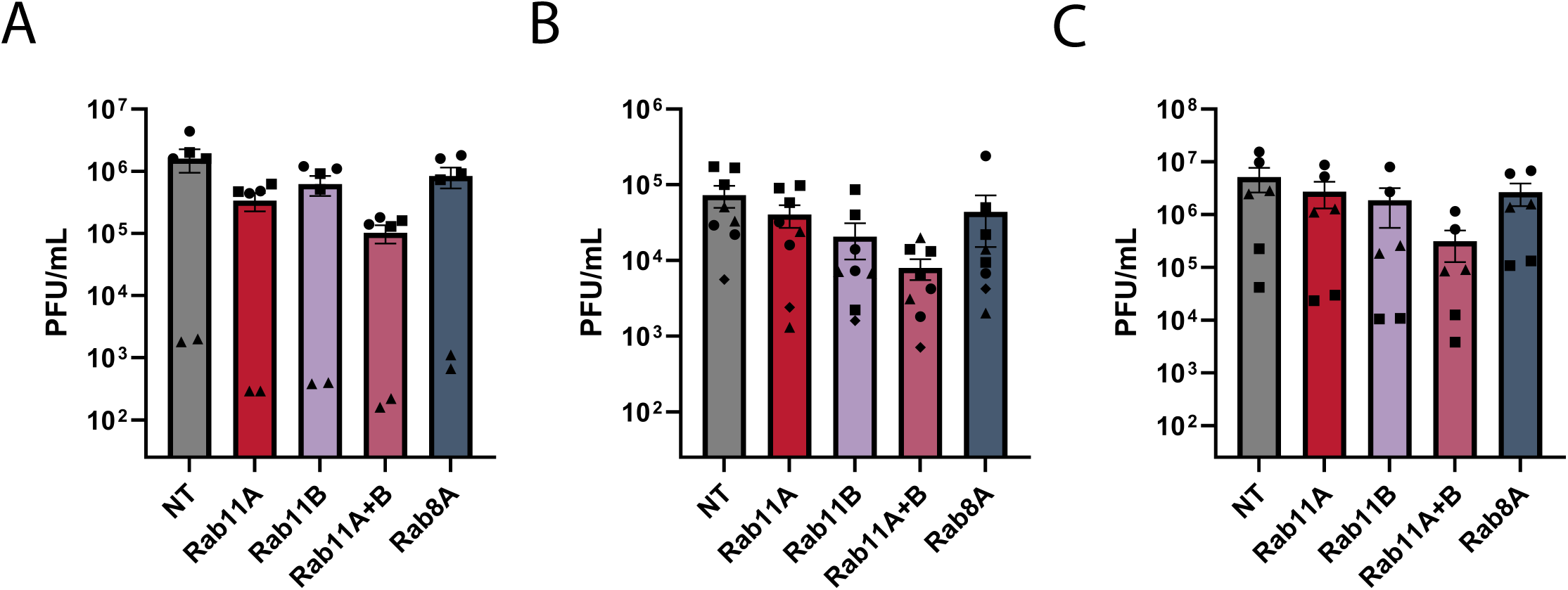
Raw titers of viral production in cells infected with recent H1N1 and H3N2 influenza A isolates show Rab11A and Rab11B are consistently required for infectious virus production in both subtypes. Raw titer data from cells infected with **A)** UVM-0478 (H1N1) [Figure 1B), **B)** UVM-1927 (H3N2) [Figure 1C], or **C)** JH-0586 (H3N2) [Figure 2A] is presented here as PFU/mL. Biological replicates are distinguished by shape (▪, ♦, ▴, or ●) with N=6 from three biological experiments (A, C) and N=8 from four biological experiments (B).

**Figure S2.**
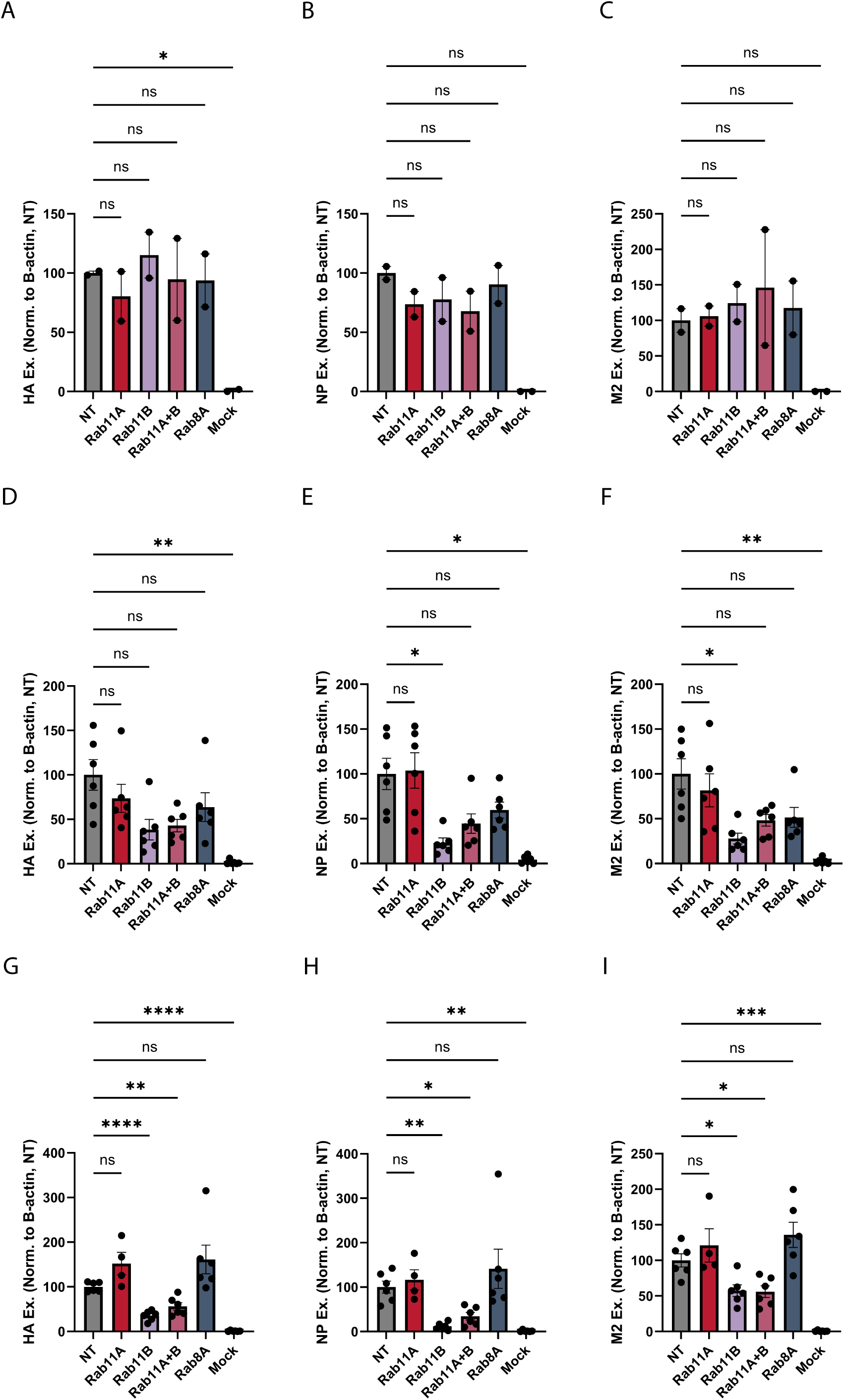
The dependence of H3N2 viral protein production in cells lacking Rab11B is seen with normalization to multiple cellular housekeeping genes. Data presented in Figure 2 and Figure 3 was alternatively quantified here by normalizing to β-actin and the average of NT control for each biological replicate; **(A)** Figure 2B, **(B)** Figure 2C, **(C)** Figure 2D, **(D)** Figure 2F, **(E)** Figure 2G, **(F)** Figure 2H, **(G)** Figure 3C, **(H)** Figure 3D, and **(I)** Figure 3E is shown for HA0 **(A, D, G)**, NP **(B, E, H)** and M2 **(C, F, I)**. Mean +/- SEM is plotted, N=2 from one biological experiment **(A-C)** and N=6 from three biological experiments **(D-I).** Statistical comparisons done using Welch’s one way ANOVA with Dunnett’s multiple comparisons [*(p<0.05), **(p<0.01), ***(p<0.001), ****(p<0.0001)].

**Figure S3.**
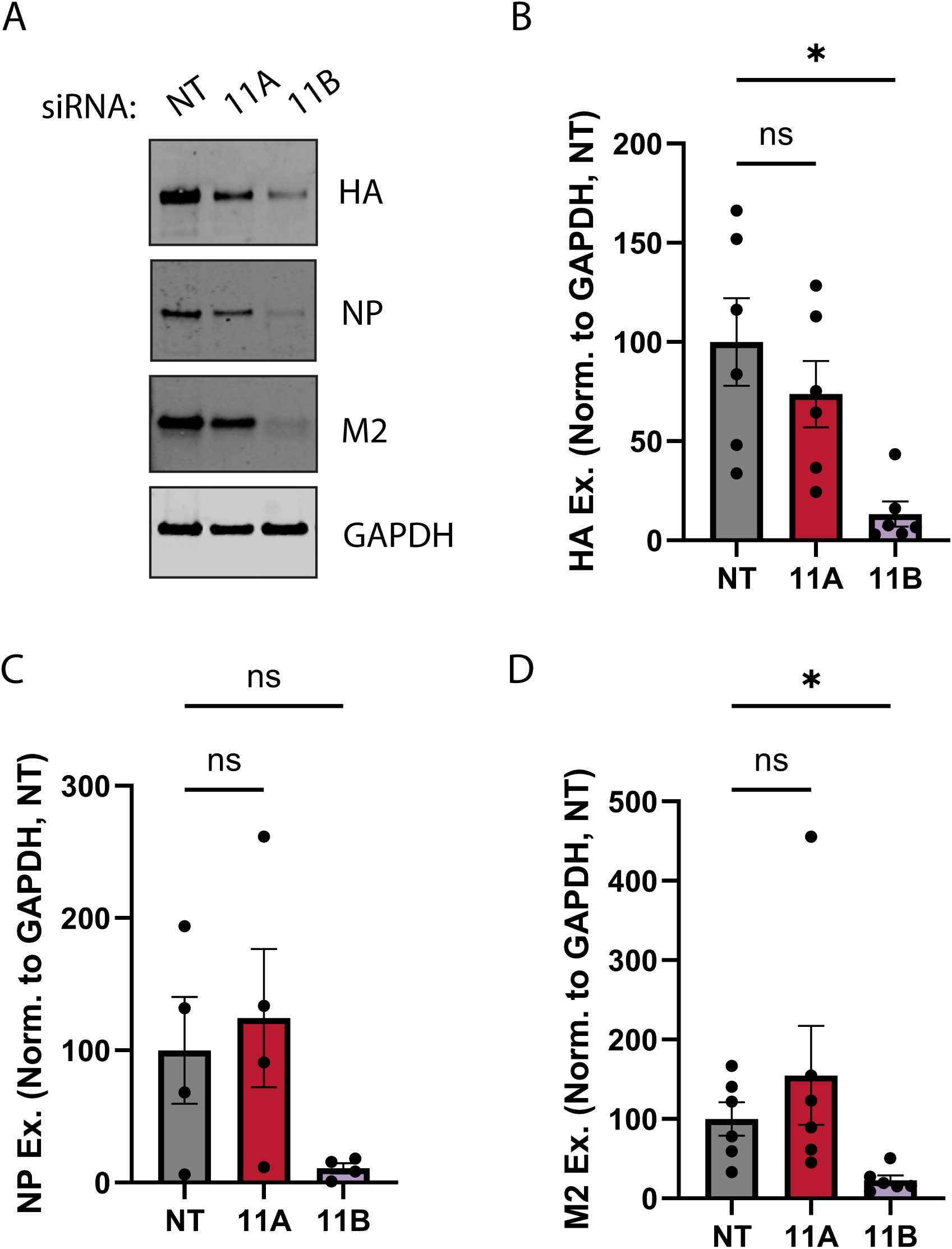
Rescue of Rab11B-dependent defect in H3N2 viral protein production in cells simultaneously depleted of Rab11A and Rab11B is not an artifact of knockdown efficiency. A549 cells were treated with siRNAs targeting Rab11A, Rab11B or a non-targeting control at half the amount typically used (5nm instead of the standard 10 nm). 48 hpt cells were infected with UVM-1927 at an MOI of 1, or mock infected. **A)** 16 hpi cell lysates were collected and proteins visualized by SDS-PAGE and western blot using rabbit anti-HA and anti-GAPDH antibodies in addition to mouse anti-NP and anti-M2 antibodies. Expression of viral proteins was quantified and normalized to GAPDH levels and the average of the NT control for each biological replicate is shown for HA0 **(B)**, NP **(C)** and M2 **(D)**. Mean +/- SEM is plotted, N=6 from three biological experiments. Statistical comparisons done using Welch’s one way ANOVA with Dunnett’s multiple comparisons (*=p<0.05).

## Notes

https://www.ncbi.nlm.nih.gov/bioproject/PRJNA1254704

### Competing Interest Statement

The authors have declared no competing interest.

### Summary of Updates

This manuscript has been revised in accordance with peer review feedback provided through Review Commons. There are now two first authors, and we have added additional experimental work to increase rigor and further advance our mechanistic understanding of how Rab11B contributes to H3N2 infection.

## REFERENCES

1. Influenza (seasonal). https://www.who.int/news-room/fact-sheets/detail/influenza-(seasonal).

2. CDC. First H5 Bird Flu Death Reported in United States. *CDC Newsroom* https://www.cdc.gov/media/releases/2025/m0106-h5-birdflu-death.html (2025).

3. Hutchinson, E. C. & Yamauchi, Y. Understanding Influenza. in Methods in Molecular Biology 1–21 (Springer New York, 2018). doi:10.1007/978-1-4939-8678-1_1.

4. Krammer, F. et al. Influenza. Nat Rev Dis Primers 4, 1–21 (2018).

5. Dourmashkin, R. R. & Tyrrell, D. A. J. Electron Microscopic Observations on the Entry of Influenza Virus into Susceptible Cells. Journal of General Virology 24, 129–141 (1974).

6. Gottschalk, A. On the Mechanism Underlying Initiation of Influenza Virus Infection. in Ergebnisse der Mikrobiologie Immunitätsforschung und Experimentellen Therapie (eds Kikuth, W., Meyer, K. F., Nauck, E. G., Pappenheimer, A. M. & Tomcsik, J.) 1–22 (Springer Berlin Heidelberg, Berlin, Heidelberg, 1959).

7. Luo, M. Influenza Virus Entry. in Viral Molecular Machines (eds Rossmann, M. G. & Rao, V. B.) 201–221 (Springer US, Boston, MA, 2012). doi:10.1007/978-1-4614-0980-9_9.

8. Fujioka, Y. et al. A Sialylated Voltage-Dependent Ca2+ Channel Binds Hemagglutinin and Mediates Influenza A Virus Entry into Mammalian Cells. Cell Host & Microbe 23, 809–818.e5 (2018).

9. Eierhoff, T., Hrincius, E. R., Rescher, U., Ludwig, S. & Ehrhardt, C. The epidermal growth factor receptor (EGFR) promotes uptake of influenza A viruses (IAV) into host cells. PLoS Pathog 6, e1001099 (2010).

10. Hale, B. G., Jackson, D., Chen, Y.-H., Lamb, R. A. & Randall, R. E. Influenza A virus NS1 protein binds p85β and activates phosphatidylinositol-3-kinase signaling. Proceedings of the National Academy of Sciences 103, 14194–14199 (2006).

11. Kumar, N., Liang, Y., Parslow, T. G. & Liang, Y. Receptor Tyrosine Kinase Inhibitors Block Multiple Steps of Influenza A Virus Replication. Journal of Virology 85, 2818–2827 (2011).

12. Kumar, N., Sharma, N. R., Ly, H., Parslow, T. G. & Liang, Y. Receptor Tyrosine Kinase Inhibitors That Block Replication of Influenza A and Other Viruses. Antimicrobial Agents and Chemotherapy 55, 5553–5559 (2011).

13. Pleschka, S. et al. Influenza virus propagation is impaired by inhibition of the Raf/MEK/ERK signalling cascade. Nat Cell Biol 3, 301–305 (2001).

14. Sieben, C., Sezgin, E., Eggeling, C. & Manley, S. Influenza A viruses use multivalent sialic acid clusters for cell binding and receptor activation. PLOS Pathogens 16, e1008656 (2020).

15. Zhu, L., Ly, H. & Liang, Y. PLC-γ1 signaling plays a subtype-specific role in postbinding cell entry of influenza A virus. J Virol 88, 417–424 (2014).

16. Peng, W. et al. Recent H3N2 Viruses Have Evolved Specificity for Extended, Branched Human-type Receptors, Conferring Potential for Increased Avidity. Cell Host & Microbe 21, 23–34 (2017).

17. Lin, Y. P. et al. Evolution of the receptor binding properties of the influenza A(H3N2) hemagglutinin. Proceedings of the National Academy of Sciences 109, 21474–21479 (2012).

18. Karakus, U., Pohl, M. O. & Stertz, S. Breaking the Convention: Sialoglycan Variants, Coreceptors, and Alternative Receptors for Influenza A Virus Entry. Journal of Virology 94, 10.1128/jvi.01357-19 (2020).

19. Sempere Borau, M. & Stertz, S. Entry of influenza A virus into host cells - recent progress and remaining challenges. Curr Opin Virol 48, 23–29 (2021).

20. Daniels, R. S., Douglas, A. R., Skehel, J. J. & Wiley, D. C. Analyses of the Antigenicity of Influenza Haemagglutinin at the pH Optimum for Virus-mediated Membrane Fusion. Journal of General Virology 64, 1657–1662 (1983).

21. Matlin, K. S., Reggio, H., Helenius, A. & Simons, K. Infectious entry pathway of influenza virus in a canine kidney cell line. Journal of Cell Biology 91, 601–613 (1981).

22. Skehel, J. J. & Wiley, D. C. Receptor Binding and Membrane Fusion in Virus Entry: The Influenza Hemagglutinin. Annual Review of Biochemistry 69, 531–569 (2000).

23. White, J., Kartenbeck, J. & Helenius, A. Membrane fusion activity of influenza virus. The EMBO Journal 1, 217–222 (1982).

24. Helenius, A. Unpacking the incoming influenza virus. Cell 69, 577–578 (1992).

25. Hutchinson, E. C. & Fodor, E. Transport of the Influenza Virus Genome from Nucleus to Nucleus. Viruses 5, 2424–2446 (2013).

26. Amorim, M. J. et al. A Rab11- and Microtubule-Dependent Mechanism for Cytoplasmic Transport of Influenza A Virus Viral RNA. Journal of Virology 85, 4143–4156 (2011).

27. Eisfeld, A. J., Kawakami, E., Watanabe, T., Neumann, G. & Kawaoka, Y. RAB11A Is Essential for Transport of the Influenza Virus Genome to the Plasma Membrane. Journal of Virology 85, 6117–6126 (2011).

28. Momose, F. et al. Apical Transport of Influenza A Virus Ribonucleoprotein Requires Rab11-positive Recycling Endosome. PLOS ONE 6, e21123 (2011).

29. Stenmark, H. Rab GTPases as coordinators of vesicle traffic. Nat Rev Mol Cell Biol 10, 513–525 (2009).

30. Veler, H. et al. The C-Terminal Domains of the PB2 Subunit of the Influenza A Virus RNA Polymerase Directly Interact with Cellular GTPase Rab11a. Journal of Virology 96, e01979–21 (2022).

31. Bhagwat, A. R. et al. Quantitative live cell imaging reveals influenza virus manipulation of Rab11A transport through reduced dynein association. Nat Commun 11, 23 (2020).

32. de Castro Martin, I. F., et al. Influenza virus genome reaches the plasma membrane via a modified endoplasmic reticulum and Rab11-dependent vesicles. Nat Commun 8, 1396 (2017).

33. Vale-Costa, S. et al. Influenza A virus ribonucleoproteins modulate host recycling by competing with Rab11 effectors. Journal of Cell Science 129, 1697–1710 (2016).

34. Han, J. et al. Host factor Rab11a is critical for efficient assembly of influenza A virus genomic segments. PLOS Pathogens 17, e1009517 (2021).

35. Vale-Costa, S. et al. ATG9A regulates the dissociation of recycling endosomes from microtubules to form liquid influenza A virus inclusions. PLOS Biology 21, e3002290 (2023).

36. Vale-Costa, S. & and Amorim, M. J. Clustering of Rab11 vesicles in influenza A virus infected cells creates hotspots containing the 8 viral ribonucleoproteins. Small GTPases 8, 71–77 (2017).

37. Nturibi, E., Bhagwat, A. R., Coburn, S., Myerburg, M. M. & Lakdawala, S. S. Intracellular Colocalization of Influenza Viral RNA and Rab11A Is Dependent upon Microtubule Filaments. J Virol 91, e01179–17 (2017).

38. Lakdawala, S. S. et al. Influenza A Virus Assembly Intermediates Fuse in the Cytoplasm. PLoS Pathog 10, e1003971 (2014).

39. Chou, Y. et al. Colocalization of Different Influenza Viral RNA Segments in the Cytoplasm before Viral Budding as Shown by Single-molecule Sensitivity FISH Analysis. PLOS Pathogens 9, e1003358 (2013).

40. Ganti, K., Han, J., Manicassamy, B. & Lowen, A. C. Rab11a mediates cell-cell spread and reassortment of influenza A virus genomes via tunneling nanotubes. PLoS Pathog 17, e1009321 (2021).

41. Bruce, E. A., Digard, P. & Stuart, A. D. The Rab11 pathway is required for influenza A virus budding and filament formation. J Virol 84, 5848–5859 (2010).

42. Ferro, E., Bosia, C. & Campa, C. C. RAB11-Mediated Trafficking and Human Cancers: An Updated Review. Biology (Basel) 10, 26 (2021).

43. Grimsey, N. J., Coronel, L. J., Cordova, I. C. & Trejo, J. Recycling and Endosomal Sorting of Protease-activated Receptor-1 Is Distinctly Regulated by Rab11A and Rab11B Proteins*. Journal of Biological Chemistry 291, 2223–2236 (2016).

44. Peterl, S. et al. Morphology-dependent entry kinetics and spread of influenza A virus. 2024.08.01.605992 Preprint at 10.1101/2024.08.01.605992 (2024).

45. Pinto, R. M., Lycett, S., Gaunt, E. & Digard, P. Accessory Gene Products of Influenza A Virus. Cold Spring Harb Perspect Med 11, a038380 (2021).

46. Petersen, J. et al. The Major Cellular Sterol Regulatory Pathway Is Required for Andes Virus Infection. PLOS Pathogens 10, e1003911 (2014).

47. Kelly, E. E., Horgan, C. P. & McCaffrey, M. W. Rab11 proteins in health and disease. Biochemical Society Transactions 40, 1360–1367 (2012).

48. Haugsten, E. M., Brech, A., Liestøl, K., Norman, J. C. & Wesche, J. Photoactivation Approaches Reveal a Role for Rab11 in FGFR4 Recycling and Signalling. TraPic 15, 665–683 (2014).

49. Silvis, M. R. et al. Rab11b Regulates the Apical Recycling of the Cystic Fibrosis Transmembrane Conductance Regulator in Polarized Intestinal Epithelial Cells. MBoC 20, 2337–2350 (2009).

50. Scapin, S. M. N. et al. The crystal structure of the small GTPase Rab11b reveals critical differences relative to the Rab11a isoform. Journal of Structural Biology 154, 260–268 (2006).

51. Chandrasekaran, A. et al. Glycan topology determines human adaptation of avian H5N1 virus hemagglutinin. Nat Biotechnol 26, 107–113 (2008).

52. Eisen, M. B., Sabesan, S., Skehel, J. J. & Wiley, D. C. Binding of the Influenza A Virus to Cell-Surface Receptors: Structures of Five Hemagglutinin–Sialyloligosaccharide Complexes Determined by X-Ray Crystallography. Virology 232, 19–31 (1997).

53. Gamblin, S. J. et al. The Structure and Receptor Binding Properties of the 1918 Influenza Hemagglutinin. Science 303, 1838–1842 (2004).

54. Ha, Y., Stevens, D. J., Skehel, J. J. & Wiley, D. C. X-ray structures of H5 avian and H9 swine influenza virus hemagglutinins bound to avian and human receptor analogs. Proceedings of the National Academy of Sciences 98, 11181–11186 (2001).

55. Ha, Y., Stevens, D. J., Skehel, J. J. & Wiley, D. C. X-ray structure of the hemagglutinin of a potential H3 avian progenitor of the 1968 Hong Kong pandemic influenza virus⋆. Virology 309, 209–218 (2003).

56. Rogers, G. N. & Paulson, J. C. Receptor determinants of human and animal influenza virus isolates: Differences in receptor specificity of the H3 hemagglutinin based on species of origin. Virology 127, 361–373 (1983).

57. Russell, R. J. et al. Structure of influenza hemagglutinin in complex with an inhibitor of membrane fusion. Proceedings of the National Academy of Sciences 105, 17736–17741 (2008).

58. Stevens, J. et al. Glycan Microarray Analysis of the Hemagglutinins from Modern and Pandemic Influenza Viruses Reveals Different Receptor Specificities. Journal of Molecular Biology 355, 1143–1155 (2006).

59. Byrd-Leotis, L. et al. Shotgun glycomics of pig lung identifies natural endogenous receptors for influenza viruses. Proc Natl Acad Sci U S A 111, E2241–2250 (2014).

60. Stevens, J. et al. Receptor specificity of influenza A H3N2 viruses isolated in mammalian cells and embryonated chicken eggs. J Virol 84, 8287–8299 (2010).

61. Byrd-Leotis, L. et al. Influenza binds phosphorylated glycans from human lung. Science Advances 5, eaav2554 (2019).

62. Sun, X. et al. Bat-Derived Influenza Hemagglutinin H17 Does Not Bind Canonical Avian or Human Receptors and Most Likely Uses a Unique Entry Mechanism. Cell Reports 3, 769–778 (2013).

63. Tong, S. et al. New World Bats Harbor Diverse Influenza A Viruses. PLOS Pathogens 9, e1003657 (2013).

64. Zhu, X. et al. Hemagglutinin homologue from H17N10 bat influenza virus exhibits divergent receptor-binding and pH-dependent fusion activities. Proceedings of the National Academy of Sciences 110, 1458–1463 (2013).

65. Karakus, U. et al. MHC class II proteins mediate cross-species entry of bat influenza viruses. Nature 567, 109–112 (2019).

66. Karakus, U. et al. MHC class II proteins mediate sialic acid independent entry of human and avian H2N2 influenza A viruses. Nat Microbiol 9, 2626–2641 (2024).

67. Chan, C.-M. et al. Hemagglutinin of influenza A virus binds specifically to cell surface nucleolin and plays a role in virus internalization. Virology 494, 78–88 (2016).

68. Achdout, H. et al. Enhanced Recognition of Human NK Receptors After Influenza Virus Infection1. The Journal of Immunology 171, 915–923 (2003).

69. Ho, J. W. et al. H5-Type Influenza Virus Hemagglutinin Is Functionally Recognized by the Natural Killer-Activating Receptor NKp44. Journal of Virology 82, 2028–2032 (2008).

70. Mandelboim, O. et al. Recognition of haemagglutinins on virus-infected cells by NKp46 activates lysis by human NK cells. Nature 409, 1055–1060 (2001).

71. Lee, J. M. et al. Deep mutational scanning of hemagglutinin helps predict evolutionary fates of human H3N2 influenza variants. Proc Natl Acad Sci U S A 115, E8276–E8285 (2018).

72. Zhou, B. & Wentworth, D. E. Influenza A Virus Molecular Virology Techniques. in Influenza Virus: Methods and Protocols (eds Kawaoka, Y. & Neumann, G.) 175–192 (Humana Press, Totowa, NJ, 2012). doi:10.1007/978-1-61779-621-0_11.

73. Iverson, T. WalkerCreek: A Nextflow Pipeline for Influenza Genome Analysis. UPHL-BioNGS (2023).

74. Ewels, P. A. et al. The nf-core framework for community-curated bioinformatics pipelines. Nat Biotechnol 38, 276–278 (2020).

75. Di Tommaso, P. et al. Nextflow enables reproducible computational workflows. Nat Biotechnol 35, 316–319 (2017).

76. Katoh, K., Misawa, K., Kuma, K. & Miyata, T. MAFFT: a novel method for rapid multiple sequence alignment based on fast Fourier transform. Nucleic Acids Research 30, 3059–3066 (2002).

77. Kurtzer, G. M., Sochat, V. & Bauer, M. W. Singularity: Scientific containers for mobility of compute. PLOS ONE 12, e0177459 (2017).

78. Minh, B. Q., et al. IQ-TREE 2: New Models and Efficient Methods for Phylogenetic Inference in the Genomic Era. Molecular Biology and Evolution 37, 1530–1534 (2020).

79. Shepard, S. S., et al. Viral deep sequencing needs an adaptive approach: IRMA, the iterative refinement meta-assembler. BMC Genomics 17, 708 (2016).

80. Chang, W. shiny: Web Application Framework for R. R package version 1.12.0. (2025).

81. Posit team. RStudio: Integrated Development Environment for R. (Posit Software, PBC, Boston, MA, 2023).

82. Nagy, A., et al. A universal RT-qPCR assay for “One Health” detection of influenza A viruses. PLOS ONE 16, e0244669 (2021).

